# Mechanism-guided mutagenesis of Rft1 to test its role as a dolichol-linked oligosaccharide scramblase in cells

**DOI:** 10.64898/2025.12.07.692794

**Authors:** George N. Chiduza, Kentaro Sakata, Hannah G. Wolfe, Faria Noor, Anant K. Menon

## Abstract

The membrane protein Rft1 is proposed to play an essential role in yeast and human cells by scrambling the glycolipid Man5GlcNAc2-PP-dolichol (M5-DLO) across the endoplasmic reticulum (ER) for protein *N*-glycosylation. While this activity has been demonstrated in liposomes reconstituted with purified Rft1, biochemical evidence of additional M5-DLO scramblases and the viability of Rft1-null *Trypanosoma brucei* suggest that scrambling may be a moonlighting function of Rft1 rather than its essential cellular role. To investigate this paradox, we used AlphaFold3 and Chai-1 to model the conformational dynamics of yeast Rft1-M5-DLO complexes. The models suggest an alternating access mechanism, typical of Multidrug/Oligosaccharidyl-lipid/Polysaccharide (MOP) superfamily transporters, in which a cationic central cavity coordinates the anionic headgroup of M5-DLO, while the dolichol tail of the lipid is accommodated through a lateral portal formed by two transmembrane helices. We used the models to design mutations to disrupt the interaction between Rft1 and the M5-DLO headgroup, and to engineer a salt bridge to block the portal and stall transport. Using a Tet-off yeast reporter strain, we tested 26 central cavity mutants and identified two that supported cell growth poorly despite being well-expressed. Strikingly, the portal-blocking mutant which lacks scramblase activity supported robust growth. These data suggest that while M5-DLO binding is important for Rft1’s essential function, scrambling activity is dispensable. We propose that Rft1’s essential role may be as an M5-DLO chaperone, capturing and routing M5-DLO propitiously on the cytoplasmic side of the ER to coordinate DLO biosynthesis.

**Importance:** Cell surface and secreted proteins are decorated with sugar chains. These chains are first assembled on a lipid carrier. Initial stages of assembly occur on the cytoplasmic side of a subcellular structure called the endoplasmic reticulum (ER). To complete assembly, the partially assembled lipid-linked sugar chain must be flipped across the ER. Here we use computationally guided cell-based assays to examine the role of the Rft1 protein in this process.

## Introduction

Rft1 is an endoplasmic reticulum (ER) membrane protein^1^, essential for the viability of yeast and human cells^2,3^. Its deficiency underlies a congenital disorder of glycosylation, RFT1-CDG^2,4,5^, associated with defects in protein *N*-glycosylation in the ER. When the level of cellular Rft1 is lowered by transcriptional repression^1,3,6,7^ or if cells express a hypomorphic variant of the protein^8^, *N*-glycosylation is compromised^3,8^, the unfolded protein response is activated^8^, and the glycolipid Man_5_GlcNAc_2_-PP-dolichol (termed M5-dolichol-linked oligosaccharide, or M5-DLO) accumulates^3^. M5-DLO is a key intermediate in the dolichol pathway that generates Glc_3_Man_9_GlcNAc_2_-PP-dolichol (G3M9-DLO), the oligosaccharide donor for *N*-glycosylation in the ER lumen (Fig. 1A)^9–13^. It is synthesized on the cytoplasmic face of the ER and must be flipped across the membrane to the luminal side to be converted to G3M9-DLO (Fig. 1A). Translocation of M5-DLO (or, less optimally, other DLOs) across the ER membrane is essential for *N*-glycosylation, and it was proposed that this key step is facilitated by Rft1^3,14^. In support of this proposal a recent report indicated that purified yeast and human Rft1 proteins are indeed capable of translocating M5-DLO across the membrane when reconstituted into synthetic lipid vesicles^15^. Because translocation in the reconstituted system occurred in the absence of metabolic energy, Rft1 can be considered to be a scramblase-class lipid transporter^16,17^.

**Figure 1.**
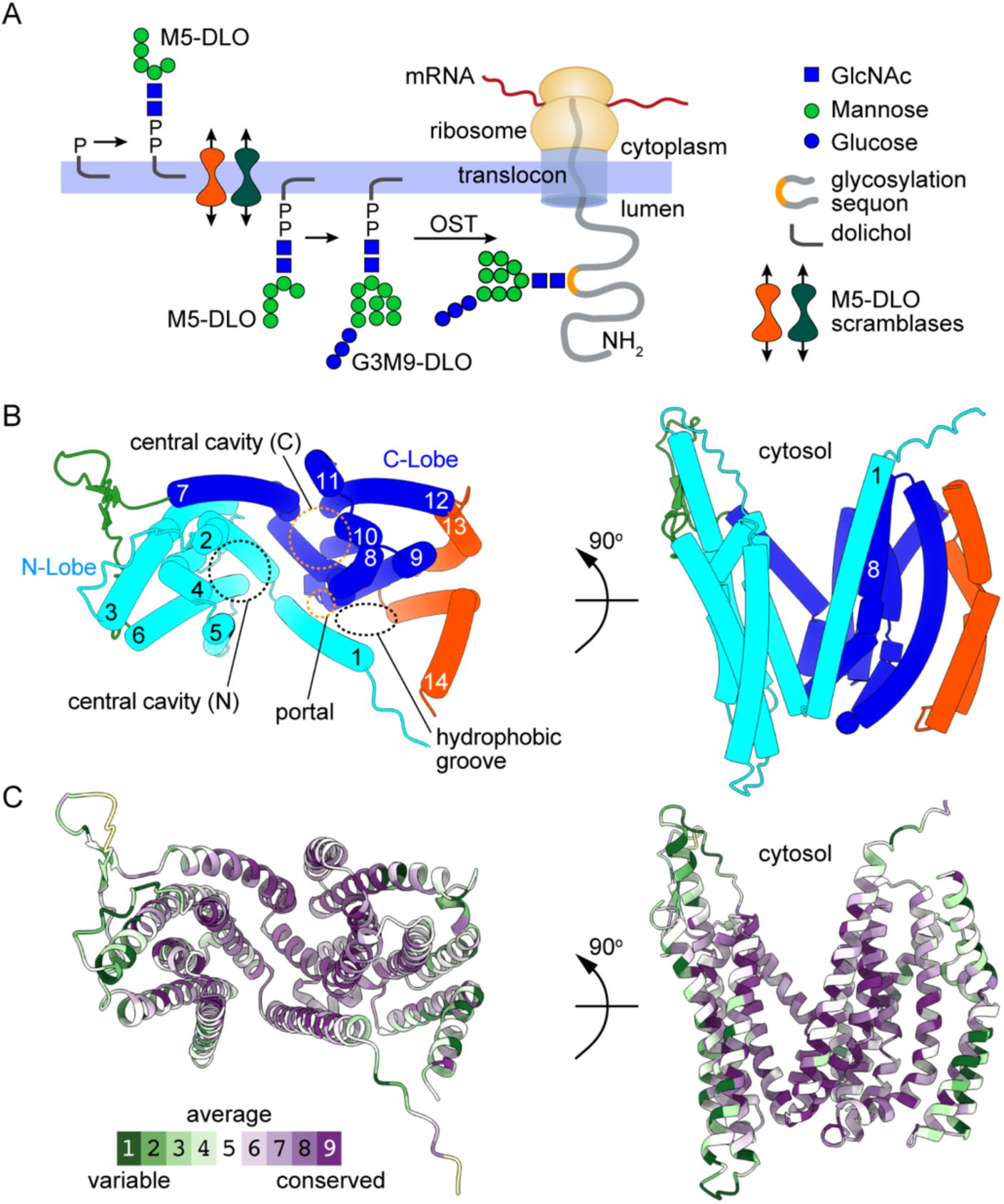
Rft1 is a MOP family transporter with homology to the lipid II transporter MurJ. **A.** Pathway of *N*-glycosylation in the ER. Dolichyl-P (top left) is converted to Man_5_GlcNAc_2_-PP-dolichol (M5-DLO) on the cytoplasmic face. M5-DLO is then transported across the membrane by redundant scramblases and converted to Glc_3_Man_9_GlcNAc_2_-PP -dolichol (G3M9-DLO). G3M9-DLO is used by oligosaccharyl-transferase (OST) on the luminal side of the ER to *N*-glycosylate nascent chains at asparagine residues within glycosylation sequons. **B.** Top and side views of the Rft1 AlphaFold model (AF-P38206-F1-v6) with helices and cavities labelled based on MurJ homology and nomenclature. Transmembrane (TM) helices comprising the N-lobe (TM1 – 6) and C-lobe (TM7 – 12) typical of MOP transporters are in cyan and blue whereas additional helices (TM13 – 14) and the long extracellular loop are in orange red and green, respectively. **C.** Alphafold model of Rft1 colored according to the calculated Consurf grade (color scale indicated).

As *N*-glycosylation is essential for cell viability, Rft1’s essentiality could be due to its ability to scramble M5-DLO across the ER membrane provided that it is the sole M5-DLO transporter in cells. However, this point is controversial. Rft1-null insect-stage cells of the early diverging eukaryote *Trypanosoma brucei* are viable and retain significant *N*-glycosylation, implying the existence of an Rft1-independent M5-DLO transport mechanism(s)^18^. Furthermore, biochemical studies show that intact microsomes prepared from Rft1-deficient yeast cells are able to convert newly synthesized GlcNAc_2_-PP-dolichol to Man_9_GlcNAc_2_-PP-dolichol and import GlcNAc_2_-PP-dolichol_15_, a water-soluble short-chain M5-DLO analog^7^. These results indicate that the microsomes have M5-DLO translocation activity despite Rft1 deficiency. In other experiments, ER membrane proteins were extracted with detergent from yeast or rat liver microsomes and reconstituted into large unilamellar vesicles. The resulting proteoliposomes were shown to translocate M5-DLO in an ATP-independent manner, indicative of the activity of a scramblase(s)^1,6,19,20^. Scramblase activity in the reconstituted system was highly specific such that higher order DLOs (Glc_1-2_Man_6-9_GlcNAc_2_-PP-dolichol) and a triantennary structural isomer of M5-DLO were scrambled much more slowly than the natural lipid^21^. However, elimination of Rft1 from the protein mixture prior to reconstitution did not have a detectable effect on activity^1,6,19^. These results indicate that the contribution of Rft1’s demonstrated scramblase activity to the overall M5-DLO scramblase activity of the ER protein mixture is minor and that at least one other M5-DLO scramblase is present in the extract.

The combined genetic and biochemical evidence suggests that our understanding of the role of Rft1 in cells is incomplete. We consider two hypotheses to resolve this seeming paradox. *First (hypothesis 1)*, Rft1 is the sole functioning M5-DLO scramblase in yeast and human, accounting for its essentiality. The additional M5-DLO scramblase(s) revealed in cell-free systems may be silent in cells. This could be for a variety of reasons. For example, the quaternary structure of the other scramblase(s), their inclusion in protein complexes^20^ or their restriction to an ER sub-domain^22^ may prevent M5-DLO access or otherwise block their activity. *Second (hypothesis 2)*, Rft1 is a dual-function ’moonlighting’ protein^23^ with a redundant scramblase activity alongside an essential function that is directly or indirectly related to DLO biosynthesis. Rft1 was previously proposed to act as a DLO chaperone^24,25^. In this scenario the protein would capture/catch M5-DLO and release it propitiously to promote its conversion to higher order DLOs. Thus, both hypotheses 1 and 2 posit that Rft1 binds M5-DLO, whereas the essential ’catch and release’ chaperone function specified in hypothesis 2 does not require Rft1 to scramble M5-DLO.

To distinguish between the two hypotheses experimentally, we used mechanism-guided mutagenesis to determine whether M5-DLO binding could be uncoupled from scrambling. As there are no experimentally determined structures of Rft1 nor any relevant data on how it might scramble lipids, we used AlphaFold3^26^ and Chai-1^27^ modeling to predict a possible mechanism and thereby to pinpoint sites critical for M5-DLO recognition and scrambling. Our efforts were guided by knowledge of mutations found in RFT1-CDG^1,2^, sequence conservation among Rft1 orthologues and the assignment of Rft1 as a member of the Multidrug/Oligosaccharidyl-lipid/Polysaccharide (MOP) transporter superfamily^28^ (Fig. S1). We specifically drew on Rft1’s relatedness to structurally characterized bacterial MOP flippases, including the Lipid II flippase MurJ^29^ and the Lipid III flippase WzxE^30^ (Table S1). Both MurJ and WzxE use an alternating access mechanism to transport cell wall lipid precursors that are structurally analogous to M5-DLO. Transbilayer transport of these lipids involves an intermediate state in which the anionic lipid is held and coordinated by essential positively charged residues within a large, cationic central cavity. The mechanistic importance of the central cavity is underscored by the fact that charge elimination or charge inversion mutations, e.g., R→A, R→D, at a number of sites within the cavity render the bacterial flippases non-functional^30–32^ (Fig. S2). Using these structural principles, we generated a series of Rft1 mutants and tested their ability to support the growth of Rft1-deficient yeast. Our results indicate that Rft1 likely operates by an alternating access mechanism, making use of a central cavity that, while relatively tolerant to mutations that would be expected to disrupt M5-DLO binding, nevertheless possesses key residues that affect function. Importantly, installation of a steric salt bridge designed to block transit of M5-DLO between the central cavity and outward-open conformation – thereby preventing scrambling – yields a protein that is able to support growth of Rft1-deficient yeast. This result aligns with hypothesis 2, i.e., that Rft1 is a dual-function ’moonlighting’ protein with a redundant scramblase activity alongside an essential M5-DLO binding function.

## Results

### Rft1 represents a distinct lineage within the MOP superfamily

A search for remote homologues using HHpred^33^ identified significant homology between Rft1 and multiple members of the MOP superfamily, including the bacterial Lipid II flippase MurJ, the Lipid III flippase WzxE, and MATE drug efflux pumps (Table S1)^28^. To define the relationship between Rft1 and bacterial lipid transporters rigorously, we reconstructed the phylogeny of the MOP superfamily using structure-guided alignments. The resulting maximum likelihood tree (Log-likelihood: -30219.220) reveals a deep evolutionary divergence between the Rft1 lineage and bacterial MOP transporters (Fig. S1). Eukaryotic Rft1 sequences form a strongly supported superclade (100% UFBoot/SH-aLRT) that is topologically distinct from the bacterial superclade. The bacterial Lipid II flippases (MurJ), including *E. coli*, *Thermosipho africanus*, and *Vibrio cholerae* orthologs, cluster into a separate, well-supported clade. Similarly, the Lipid III flippases (WzxE) and O-antigen transporters form a distinct cluster separate from both MurJ and Rft1. Within the Rft1 superclade, the topology recapitulates established organismal phylogeny: *Homo sapiens* and *Mus musculus* orthologs cluster as sister taxa with maximal support (100%), while the *Saccharomyces cerevisiae* and *Candida albicans* fungal orthologs form a distinct subclade (100% support). Crucially, Rft1 does not nest within the MurJ lineage but instead shares a common ancestor with the MurJ, Wzx, and MATE families. This topology identifies Rft1 as a distinct structural homolog within the MOP superfamily (Fig. S1).

While the canonical MOP fold (exemplified by Wzx and MATE transporters) consists of 12 transmembrane (TM) helices, the high-confidence models of Rft1 generated using AlphaFold^26^ and Chai-1^27^ predict a 14-transmembrane (TM) helix topology, organized into N-terminal (TM1–6) and C-terminal (TM7–12) lobes, followed by two additional C-terminal helices (TM13–14) (Fig. 1B, Fig. S2A). The TMs of Rft1 are predicted with high confidence (predicted local-distance difference test (pLDDT) > 70), except for TM1, the C-terminal half of TM7, and the N-terminal halves of TMs 3 and 14 (50 < pLDDT < 70), suggesting inherent flexibility in these regions (Fig. S2A)^34^. The two lobes form a central cavity connected to a membrane-facing hydrophobic groove (formed by TMs 8, 9 and 14) via a "portal" between TMs 1 and 8 (Fig. 1B). Sequence conservation analysis of 77 Rft1 orthologues revealed that residues lining the cationic central cavity are highly conserved (ConSurf grade > 5) (Fig. 1C, Fig. S3).

Strikingly, MurJ flippases possess a similar non-canonical 14-TM topology with a cationic central cavity that accommodates the anionic Lipid II head group, and a hydrophobic groove formed by the two additional helices (TM13 and TM14) on the protein’s surface, that is positioned to accommodate the long (C_55_) undecaprenyl tail during flipping^30,31^. MurJ from an *Arsenophonus* endosymbiont was the top hit in our HHpred search (99.81% probability, 11% sequence identity) (Table S1) and this structural homology is further supported by high structural similarity (C_α_ RMSD = 1.4 Å) between the predicted Rft1 structure (AlphaFold Database: AF-P38206-F1-v6) and the crystal structure of *Thermosipho africanus* MurJ in the inward-open conformation (PDB ID: 6NC7)^30^ (see below and Fig. S2B,C). Consequently, we use well-established MurJ nomenclature as a convenient structural and mechanistic yardstick to describe Rft1’s features (Fig. 1B).

### Predicting Rft1’s scramblase mechanism

MurJ transports Lipid II across the bacterial inner membrane using an alternating access mechanism involving transitions between inward-open and outward-open conformations. Transport is powered by membrane potential (ΔΨ) and coupled to Na^+^ antiport which is required to reset the transporter from the outward-open to the inward-open state^35^. Although the energy requirement makes MurJ a flippase-type transporter, distinguishing it from Rft1 which appears to be a scramblase, the pathway for Lipid II translocation within MurJ nevertheless suggests how Rft1 might transport M5-DLO.

We modeled the interaction between Rft1 and M5-DLO with AlphaFold3 and Chai-1, using the experimentally determined topology of the protein (N- and C- termini facing the cytoplasm, i.e., inward)^1^ to orient the models. Initially, with M5-DLO defined using the Simplified Molecular Input Line Entry System (SMILES) format, both tools successfully docked the substrate into the central cavity (Fig. 2). However, Chai-1 generated superior stereochemical models, recovering the canonical chair-like conformations for the mannose units of the headgroup. AlphaFold3 outperformed Chai-1 in generating realistic models of M5-DLO, when the ligand was defined using the Chemical Component Dictionary (CCD) format to define the glycolipid components linked via the bondedAtomPairs (BPA)syntax^36^. Here AlphaFold3 successfully modelled the correct topology and stereochemistry of the glycan, specifically the *N*-acetyl-α-D-glucosamine residues which had longer than ideal bond lengths in the Chai-1 models (Fig. S4). Crucially, the Chai-1 algorithm predicted partially occluded conformations for M5-DLO-bound states, whereas AlphaFold3 only modeled the inward facing open conformation. Thus, the outputs of the two algorithms are complementary.

**Figure 2.**
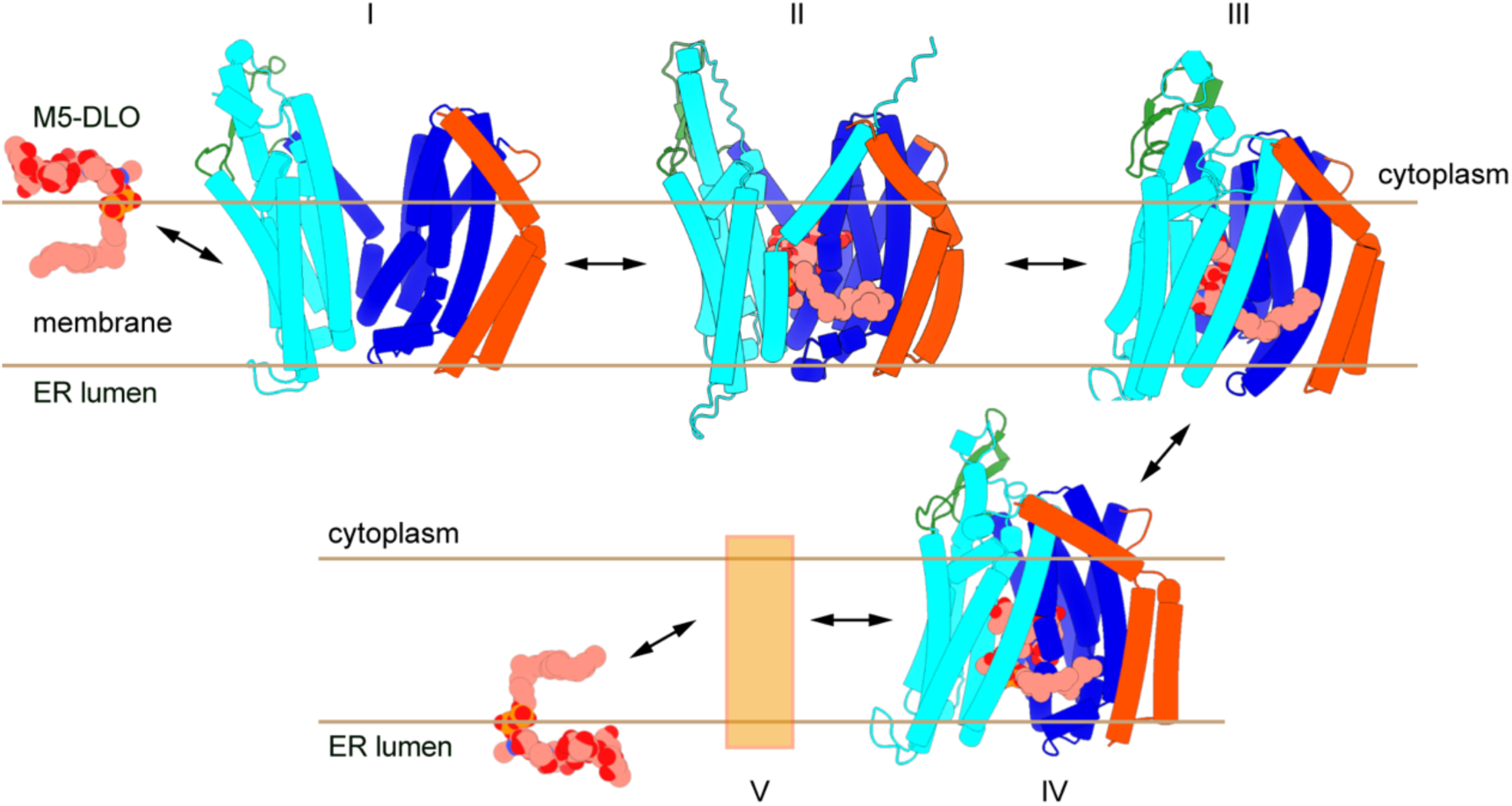
Model of transbilayer translocation of M5-DLO by Rft1. Inward open (I, II), occluded (III) and outward-facing partially occluded (IV) conformations of apo- and M5-DLO bound Rft1 predicted by Chai-1 (except for structure II predicted by AlphaFold3) and presented in the sequence of a hypothetical alternating access translocation pathway, I → V, moving M5-DLO from the cytoplasmic face of the ER to the luminal side. V is a place holder for the outward-facing open conformation (not modelled by Chai-1) needed to discharge/import M5-DLO on the luminal side. As Rft1 is a scramblase, the reverse pathway (V → I) is also expected. The structures are colored by lobe as in Fig. 1B. The M5-DLO model used has 4 isoprene units in its dolichol tail for Chai-1 and 8 isoprene units for AF3, shorter than the 14-19 isoprenes typically found in yeast dolichol (see Methods).

In the docked models (Fig. 2, Fig. 3A), the M5-DLO glycan is sequestered within the central cavity, stabilized by a network of hydrogen bonds with residues in TMs 2, 7, 8 and 10, while the diphosphate neck electrostatically engages R334 and K34 (Fig. 3B, right panel). The lipid tail traverses the lateral portal formed between TM1 and TM8 to engage the hydrophobic groove (Fig. 3). The volume of the central cavity, measured using CASTp volumetrics^37^, decreases from ∼13,000 Å³ in the inward-open state to ∼6,000 Å³ in the occluded state. MoloVol^38^ analysis reveals that this internal capacity vastly exceeds the molecular volume of the complete polar headgroup, glycan plus diphosphate, 1,067.5 Å³. Because the cavity remains cavernous even at its most constricted, occlusion effectively traps the substrate by sealing the gating portals rather than through tight steric constriction inside the chamber. Subsequent transition to the lumen-facing state expands the cavity to ∼9,000 Å³ (Fig. S5). Although this state provides immense internal volumetric clearance for the substrate to dissociate from its binding network, structural quantification confirms that the luminal gate itself remains restricted. Specifically, the central cavity mouth area of this partially occluded conformation measures ∼303 Å², compared to ∼960 Å² for the inward-open conformation (Fig. S5A & C). Thus, this state likely represents a pre-release intermediate as further expansion of the structure on the luminal gate would be needed to allow egress/entry of the substrate. As this more dilated conformation was not sampled in our analyses it is depicted as a placeholder box in Figure 2. We note that a continuous transmembrane channel is not observed in any of the structures (Fig. S5), consistent with an alternating access mechanism which distinguishes Rft1 from other scramblases that feature a transmembrane hydrophilic groove for lipid transit^39^.

**Figure 3.**
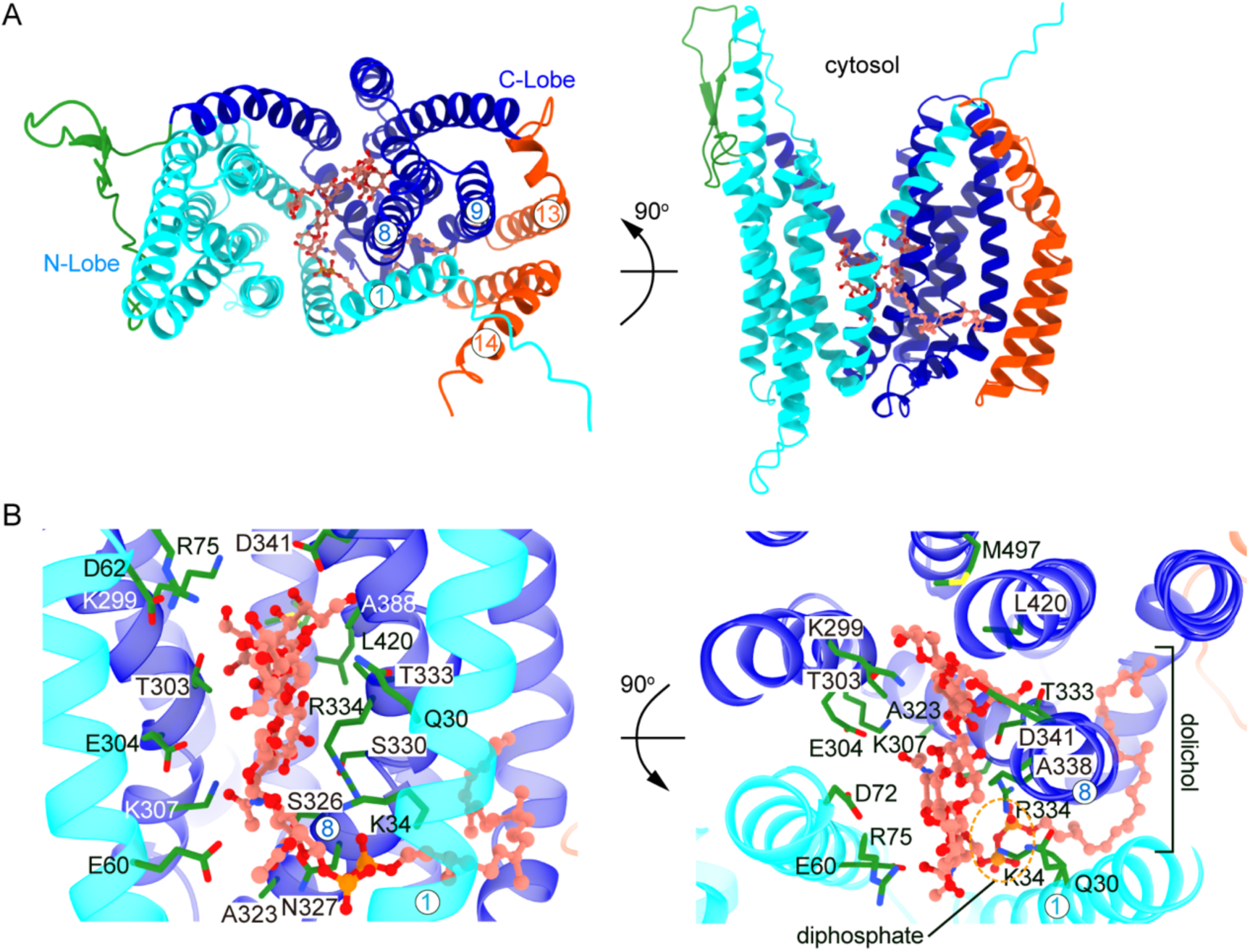
M5-DLO docked into an inward-open conformation of Rft1. **A.** Rft1-M5-DLO complex (top and side views) predicted by AlphaFold3 using CCD/BPA hybrid syntax for defining glycan structures. **B.** Chai-1 predicted pose for M5-DLO bound to Rft1 in the occluded conformation (Fig. 2, structure III), showing residues predicted to interact with M5-DLO. Atoms of M5-DLO (carbon - salmon; nitrogen - blue and phosphorus – orange) co-folded with Rft1 are depicted as spheres in panel A and sticks and balls in panel B. Certain TM helices are labeled (numbers within white circles) in the left image (A), and in both images (B).

### Mutations predicted to affect M5-DLO binding

Our models feature specific interactions between Rft1 and M5-DLO that, when disrupted, should affect binding and scrambling. We identified 64 residues within 5 Å of the docked M5-DLO molecule using the Chai-1 models; notably, five of these residues correspond to positions in human Rft1 that are mutated in RFT1-CDG, underscoring their importance (Table S2). Based on conservation (ConSurf grade ≥ 8) and contact frequency in the 5 predicted conformational poses, we selected 21 of the 64 residues for mutagenesis screening (Table 1). For 13 of these, we used charge inversion or neutralization, (e.g., K34D and R334A) for mutagenesis, based on the approach taken to analyze MurJ. A key feature of MurJ is the electrostatic environment of the central cavity, which is lined with conserved residues including a charged triad (R24 and D25 on TM1 and R255 on TM8 in *T. africanus* MurJ (in *E. coli* MurJ the third residue of the triad is R270)) which stabilizes the anionic diphosphate neck of Lipid II^29,31,32,40^ (Fig. S2C). Charge inversion or alanine substitution of these and other key central cavity residues resulted in a total loss of MurJ’s flippase activity, indicating a strict requirement for precise electrostatic coordination of the translocating substrate^29,31,32,40,41^. Thus, we anticipated that similar mutations in Rft1, e.g., at positions K34 and R334 (Fig. 3B, Table 1), would result in a non-functional protein. For the remaining residues we made polar to hydrophobic mutations, or substituted amino acids absent at the orthologous position across the 77 species analyzed (e.g., G424Q and L420Q), the rationale being that these residues are potentially underrepresented in extant species because they compromise fitness. Finally, we generated Rft1 variants carrying single, double and triple mutations at a highly conserved site (T303L, S326L, N327L)(Consurf Grade >8) predicted to mediate M5-DLO glycan recognition in all 5 Chai-1 models, Engineering a portal blockade to prevent scrambling. To distinguish between our two hypotheses, we sought to uncouple substrate binding from scrambling by engineering a portal mutant to block movement of M5-DLO during the conformational transition between the occluded and outward-facing states (structures III and IV, respectively, Fig. 2, Fig. S6). In structurally characterized MOP flippases like MurJ, transmembrane helices TM1 and TM8 form a lateral portal that connects the continuously solvated hydrophilic central cavity to the lipid bilayer^42,43^. This portal must dilate and close to facilitate the transit of M5-DLO’s lipid tail. Residues F38 and Y328 localize precisely at this dynamic, continuously hydrated protein-lipid boundary (Fig. 7A; Table1). Substituting these native hydrophobic residues with oppositely charged amino acids (D and K, respectively) introduces a structurally restrictive electrostatic tether. Our Chai-1 models indicate that the engineered D38 and K328 side chains pack in close proximity in the fully occluded conformation (Fig. 7A, helices depicted in grey). Although the unrelaxed predicted distance of 2.27 Å between D38 (OD1) and K328 (NZ) falls below the standard 2.5 to 3.0 Å van der Waals contact optimum for biomolecular salt bridges^44^, this apposition nevertheless strongly supports a highly compact interaction. Crucially, the persistent solvation of the central cavity elevates the local dielectric environment, mitigating the energetic penalty of burying charges at the membrane interface. Thus, this microenvironment stabilizes D38 and K328 in their fully ionized states, enabling the formation of a robust electrostatic salt bridge. Transitioning to the outward-facing partially occluded conformation (Fig. 2, III) requires the interatomic distance between TM1 and TM8 to increase significantly, modeled here as a dilation to 4.19 Å (Fig. 7A, colored helices). This distance is within the 4.5 Å strict cutoff for a completely unbound electrostatic interaction^44^. Because the required 4.19 Å dilation remains shorter than this critical dissociation threshold, the two residues are predicted to remain tightly coupled electrostatically, presenting a significant kinetic barrier for the TM1-TM8 portal dilation necessary to release the M5-DLO substrate on the luminal side. Consequently, the F38D/Y328K portal mutation functions as a potent lock. By tethering TM1 to TM8 through this highly stable salt bridge, the engineered interaction is proposed to physically obstruct substrate translocation and stall the alternating access cycle, restricting Rft1’s dynamics to the cytoplasmic leaflet, where it remains fully capable of substrate capture and release (Fig. S6).

**Table 1.**
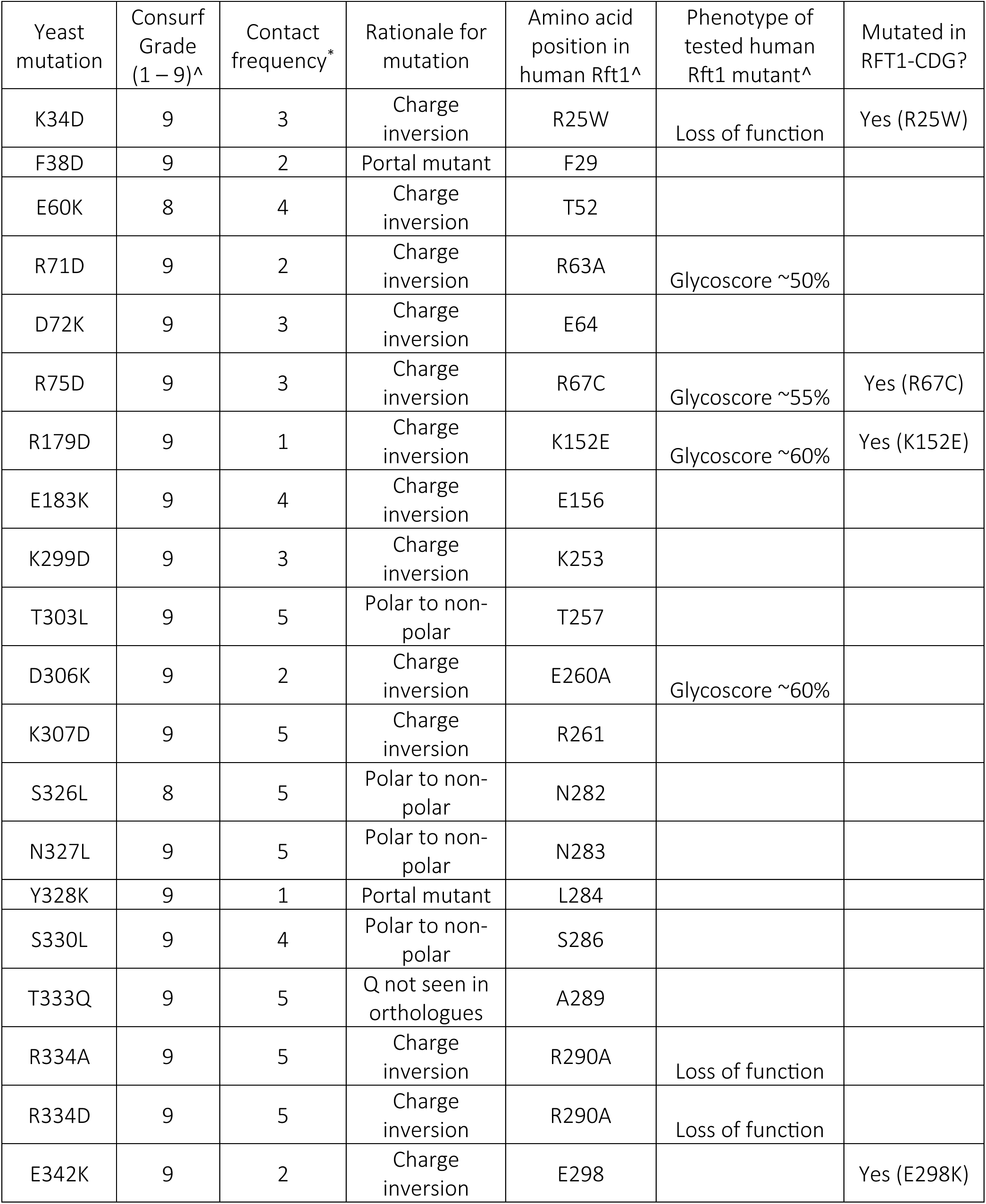
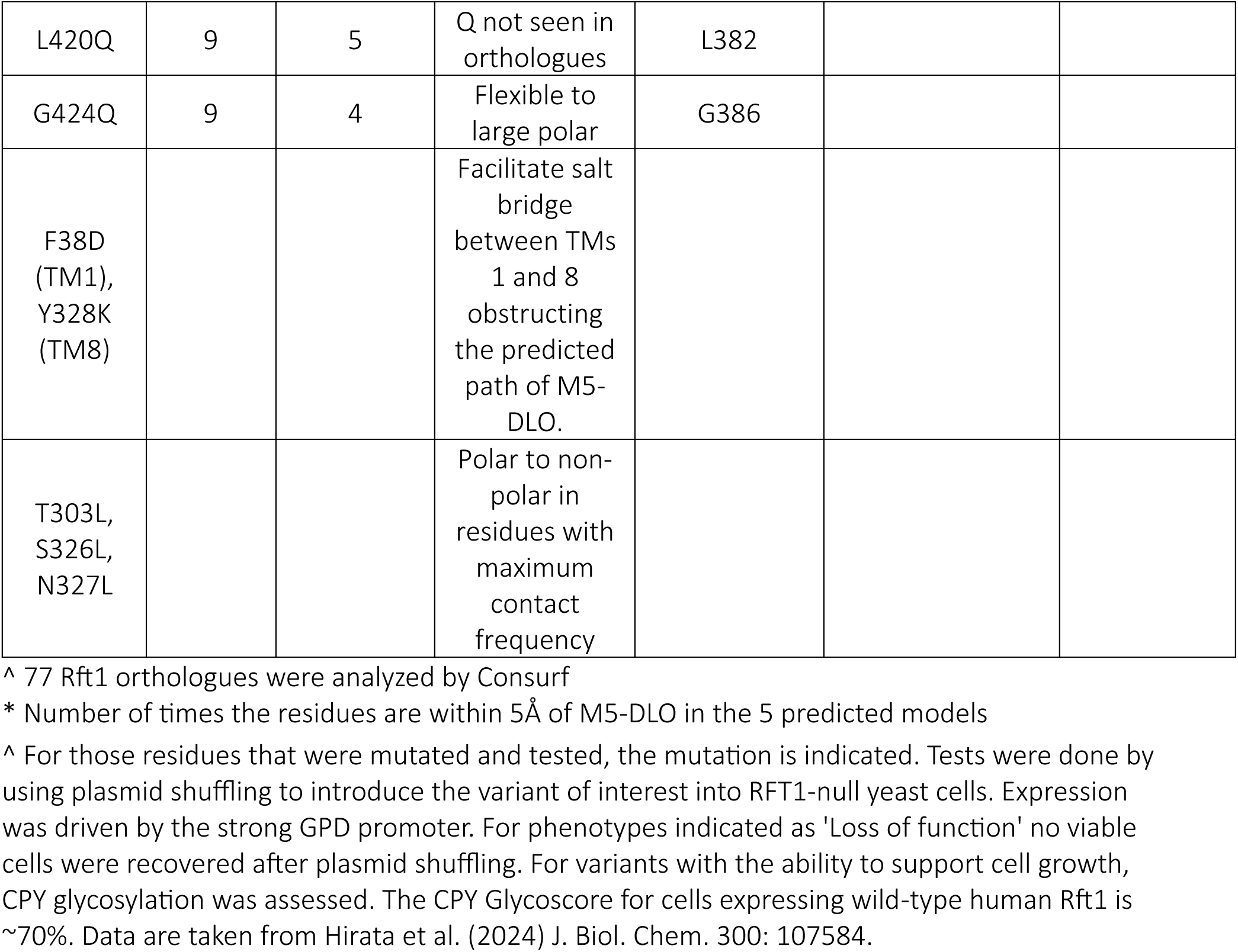
Mutations studied in this paper.

A reporter strain to test functionality of Rft1 mutants. We considered several cell-based methods to test the functionality of Rft1 mutants and opted for a promoter replacement system in which transcription of the *RFT1* gene in yeast is controlled by doxycycline (Dox)^45,46^. In these haploid Tet-off cells the addition of Dox causes rapid cessation of transcription, followed by loss of Rft1 through degradation and dilution during cell division^47^. As Rft1 is essential, addition of Dox is expected to cause the cells to grow more slowly as Rft1 levels fall.

To validate and characterize the Tet-off strain, we transformed the cells with an empty vector (EV) or a vector encoding C-terminally 3xFLAG-tagged Rft1 under control of either its own promoter (*P_RFT1_*) or the medium strength ADH promoter (*P_ADH_*)^48^. We previously established that FLAG-tagging does not interfere with the ability of Rft1 to support cell growth^1^. The transformants were serially diluted onto normal agar selection plates (Fig. 4A, left panel) as well as plates containing Dox (Fig. 4A, right panel). All strains grew on the control plates; however, only the cells with plasmid-encoded Rft1-3xFLAG grew on the +Dox plates (Fig. 4A), with the *P_RFT1_-RFT1* cells growing slightly more slowly than the *P_ADH_-RFT1* cells.

**Figure 4.**
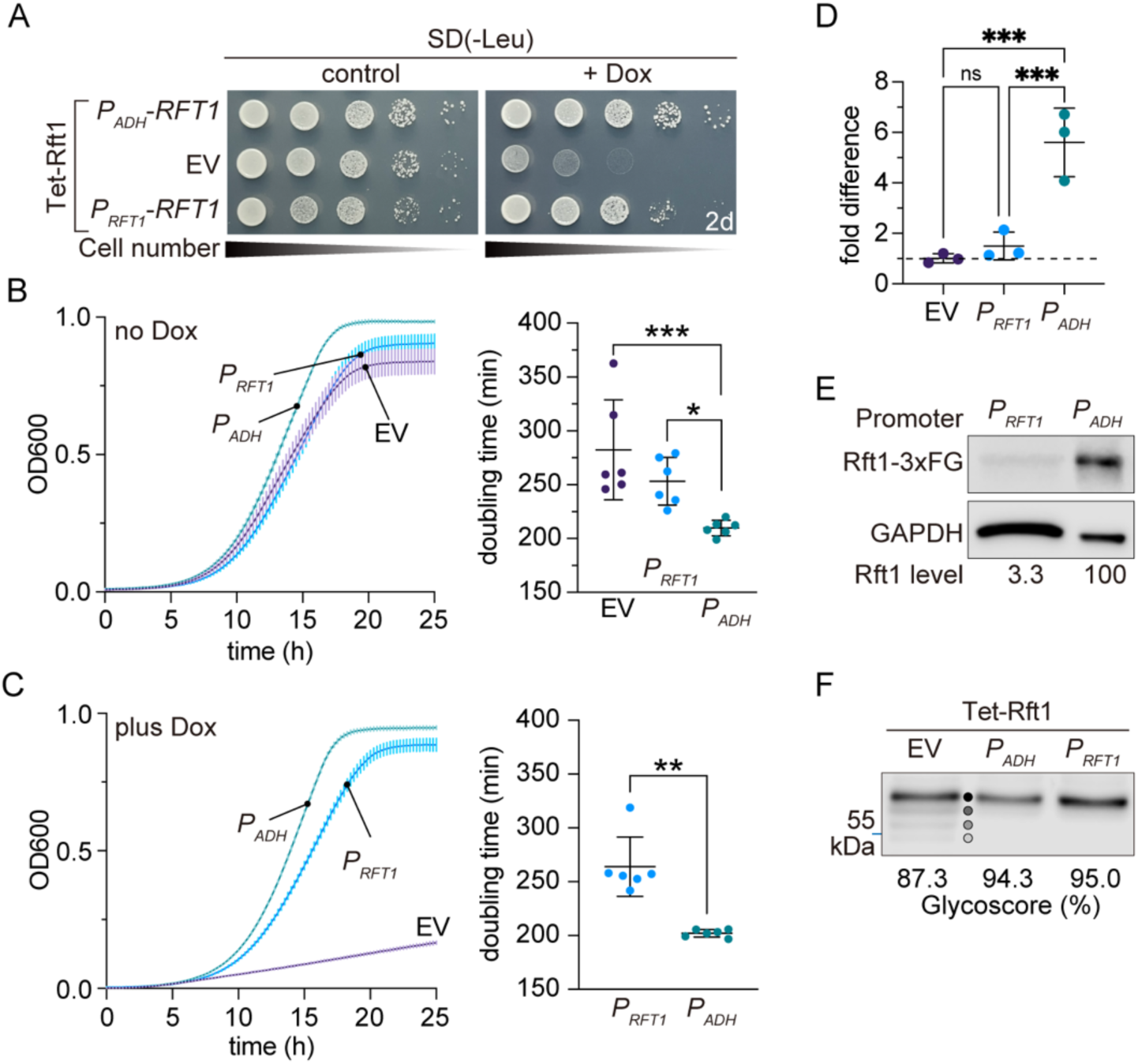
Tet-off yeast strain to assess Rft1 function. **A.** Serial 10-fold dilutions of Tet-off Rft1 cells (YAKM147) transformed with an empty vector (EV) or vectors encoding C-terminally 3xFLAG-tagged Rft1 under control of its own promoter (*P_RFT1_*) or the ADH promoter (*P_ADH_*) were spotted onto an SD(-Leu) plate (left panel) or an SD(-Leu) plate supplemented with 10 µg/mL Dox (right panel). The plates were incubated at 30°C for 2 days before being photographed. **B, C.** Growth at 30°C of suspension cultures of the same cells as in panel A. Dox (0.5 µg/mL) was added at t=0 h to the +Dox samples. Cells were initially cultured in SD(-Leu) media before being diluted into rich media (± Dox) for the growth assay. Doubling times were calculated from data in the OD600 range ≥0.5 to ≤0.65. Statistical significance in panel B was determined by ordinary one-way ANOVA (***p=0.0006, *p=0.045, comparison between EV and *P_RFT1_* data was not significant), and in panel C by unpaired t test (two-tail)(**p=0.0025). **D.** Fold-difference in *RFT1* transcript levels determined by qPCR. Statistical significance was determined by ordinary one-way ANOVA (***p=0.0006 (*P_ADH_* vs *P_RFT1_*) p=0.0003 (*P_ADH_* vs EV)). **E.** Immunoblotting to compare levels of ectopically expressed Rft1-3xFLAG in *P_RFT1_-RFT1* and *P_ADH_-RFT1* cells. 10-fold more of the *P_RFT1_-RFT1* cell extract was loaded in order to visualize Rft1-3xFLAG in the sample. The mean ratio of the FLAG signal to that of GAPDH (loading control) is indicated below the blot (Rft1 level), normalized to the value for the *P_ADH_-RFT1* cell sample (range 0.7–6.2, n=4). **F.** *N*-glycosylation of CPY. Protein extracts from EV, *P_RFT1_-RFT1* and *P_ADH_-RFT1* cells (grown without Dox) were analyzed by SDS-PAGE immunoblotting with anti-CPY antibodies. The filled circles (gradient of grey) indicate CPY glycoforms with 4. 3, 2 and 1 *N*-glycan(s). The Glycoscore was calculated from the single immunoblot shown.

We next tested the Dox sensitivity of cell growth in liquid culture using a plate reader, recording cell density (OD600) every 15 min with 5 s of orbital shaking of the multi-well plate prior to each measurement. Cells were initially cultured in selection media before being diluted into rich media (± Dox) for the growth assay. All transformants grew well in the absence of Dox, although the doubling time (mean ± S.D., n=6) of the EV cells (4.7 ± 0.78 h) and *P_RFT1_-RFT1* cells (4.2 ± 0.37) was ∼20-30% greater than that of *P_ADH_-RFT1* cells (3.5 ± 0.12 h), indicating that Rft1 expressed via the Tet promoter system or ectopically under control of its own promoter is limiting for growth (Fig. 4B). When 0.5 µg/mL Dox was included in the growth medium the doubling time of the Rft1-expressing cells did not change (the *P_ADH_-RFT1* cells grew faster than *P_RFT1_-RFT1* cells as in the minus Dox condition) but that of EV cells increased to >20 h (Fig. 4C). The Dox concentration used was adequate as the growth rate of EV-transformed cells did not slow further when a higher concentration of Dox (1.0 µg/mL) was used.

To determine the growth-limiting level of Rft1 in EV and *P_RFT1_-RFT1* cells versus *P_ADH_-RFT1* cells, we used qPCR to quantify *RFT1* transcripts in the three strains. Transcript levels in the *P_RFT1_-RFT1* cells and *P_ADH_-RFT1* cells were 1.5 and 5.6-fold higher, respectively, than in the EV cells (Fig. 4D). After correcting for mRNA transcribed from genomic *RFT1*, i.e., the level in the EV cells, this result indicates that there is a ∼10-fold difference in the level of *RFT1* mRNA produced ectopically in *P_RFT1_-RFT1* cells versus *P_ADH_-RFT1* cells, suggesting an order of magnitude difference in the level of expressed protein. We confirmed this by directly assessing relative protein levels via anti-FLAG immunoblotting. As shown in Figure 4E, the Rft1-3xFLAG signal in *P_ADH_-RFT1* cells is ∼30-fold higher than in *P_RFT1_-RFT1* cells, consistent with the mRNA data.

A meta-analysis of published datasets reported about 1000 copies of Rft1 per cell (coefficient of variation 85)^49^. We previously expressed Rft1-3xFLAG under control of the strong GPD promoter and estimated about 5,500 copies of the tagged protein per cell^1^. As expression driven by the ADH promoter is at least 5-fold lower^48^ (see Fig. 7C) we estimate at most 1000 copies of Rft1-3xFLAG in *P_ADH_-RFT1* cells and fewer than 100 copies per cell of the endogenous protein. To verify this estimate, we used anti-FLAG immunoblotting to compare Rft1-3xFLAG in the *P_ADH_-RFT1* cells with 3xFLAG-tagged bovine opsin of known concentration as previously described^1^. The measurement indicated 300 ± 78 (mean ± S.D., n=5) copies per cell, consistent with our estimate. Combined with the qPCR and expression data (Fig. 4D,E) this suggests that the level of Rft1 in the Tet-off and *P_RFT1_-RFT1* cells is very low, 20 and 30 copies per cell, respectively, far lower than indicated by the meta-analysis results^49^. Thus, Rft1’s essential function in cells is achieved by very few copies of the protein. We note that previous estimates of the abundance of M5-DLO scramblase(s) in yeast ER suggest that the scramblase(s) represent ∼1% by weight of ER proteins^19^, indicative of several thousand copies and far in excess of our estimate of the Rft1 copy number.

The level of Rft1 clearly affects growth as *P_ADH_-RFT1* cells grow ∼20% faster than *P_RFT1_-RFT1* cells (Fig. 4B,C). To learn if this effect is related to the ability of the cells to execute *N*-glycosylation we used immunoblotting to analyze the glycosylation status of carboxypeptidase Y (CPY), a vacuolar protein with 4 *N*-glycans. Interestingly, CPY was fully glycosylated in both *P_RFT1_-RFT1* and *P_ADH_-RFT1* cells grown in the absence of Dox, and somewhat hypo-glycosylated in EV cells (Fig. 4F). This result points to a possible role for Rft1 in cell growth in addition to – or instead of – *N*-glycosylation.

Our results establish the Tet-off Rft1 strain as a suitable platform to interrogate the ability of the proposed Rft1 mutants to support growth as it has the requisite Dox-sensitive growth characteristics that are responsive to ectopically expressed Rft1. The expression level of the Rft1 variants can be tuned by using promoters of different strengths (*P_RFT1_*<*P_ADH_*<*P_GPD_*) (Fig. 4D,E, see Fig. 7C), with growth being read out via spotting assays on agar plates as well as in suspension culture. Whereas the *P_ADH_* and *P_GPD_* promoters result in different degrees of over-expression, the *P_RFT1_* promoter results in ’under-expression’ as it generates 2-fold fewer copies of the wild-type protein on Dox media than would be present in EV-containing Tet-off cells grown in normal media (Fig. 4D).

Cell-based analysis of Rft1 variants with predicted M5-DLO binding defects. We transformed Tet-off cells with *P_ADH_-RFT1* plasmids encoding each of the proposed Rft1 point mutants (Table 1) with a C-terminal 3xFLAG tag. On serially diluting the cells onto plates containing Dox media, we found that 19 of the 21 single mutants supported essentially normal growth, comparable to the growth observed in cells transformed with a plasmid encoding wild-type (WT) Rft1-3xFLAG (Fig. 5A, top row) and contrasting with that of EV-transformed cells which failed to grow (Fig. 5A, second row). To complement the spotting assays, we measured growth of the transformed cells in liquid culture. All the cells grew similarly in Dox-containing media (Fig. 5B), with slight variations in doubling time (<10% higher in a few cases, when compared to cells expressing WT Rft1).

**Figure 5.**
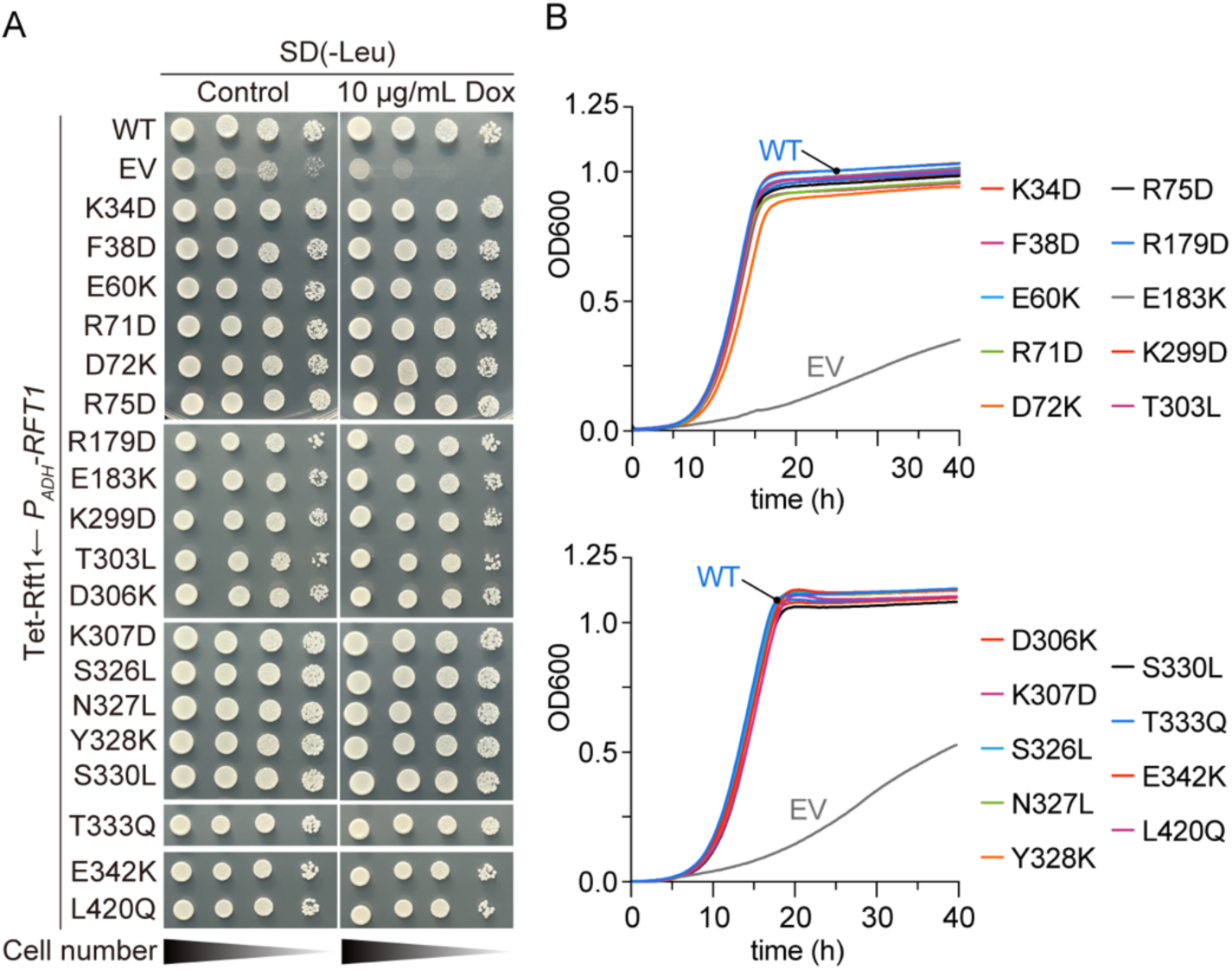
Functional test of Rft1 single mutants. **A.** Serial 10-fold dilutions of Tet-off Rft1 cells transformed with *P_ADH_-RFT1* plasmids encoding various Rft1 point mutants were spotted onto SD(−Leu) plates without (left panel) or with 10 µg/mL Dox (right panel). The plates were incubated at 30°C for 3 days before being photographed. Images from several plates are assembled in the figure – each plate had WT and EV controls as shown at the top of the panels for the first pair of plates (these controls are not shown for each plate for clarity of presentation). **B.** Growth at 30°C of the same cells as in panel A. The curves were obtained with suspension cultures in the presence of Dox as described in Fig. 4B,C. The line in each case represents the mean of 6 measurements (error bars are not shown for clarity of presentation)

Immunoblotting against the 3xFLAG epitope tag revealed 0.3-2.5-fold variation in expression level of the mutants relative to wild-type Rft1 (Fig. S7). Rft1(T303L) exhibited the lowest expression of all mutants, at ∼30% of matched wild-type levels. As this low abundance is sufficient to support robust growth, the T303L mutation appears to affect stability of the protein rather than its activity, a point further demonstrated by CPY *N*-glycosylation which was similar in Rft1(T303L)- and Rft1(WT)-expressing cells grown on Dox (Fig. S8). We used the low expression of Rft1(T303L) to identify a lower bound for the number of copies of Rft1 needed to support growth. Thus, we used the weak *P_RFT1_* promoter to express Rft1(T303L) (Fig. S9) and included Rft1(D72K) for comparison as it expresses as well as the wild-type protein (Fig. S7). As shown in Figure S9A,B, *P_RFT1_-RFT1(D72K)* transformed Tet-off cells grew well on Dox (comparable to matched Rft1(WT)-expressing cells), but *P_RFT1_-RFT1(T303L)* transformed cells grew poorly, suggesting that for the latter mutant the expression level must be close to the minimal copy number required to retain viability.

T303 is part of a cluster of highly conserved polar residues, including S326, and N327, that is predicted to engage the polar M5-DLO glycan (Fig. 6A). We found that single (Fig. 5), double and triple (Fig. 6B,C) leucine substitutions of this cluster had no effect on the ability of the resulting Rft1 variants to support growth of the Tet-off cells on Dox media.

**Figure 6.**
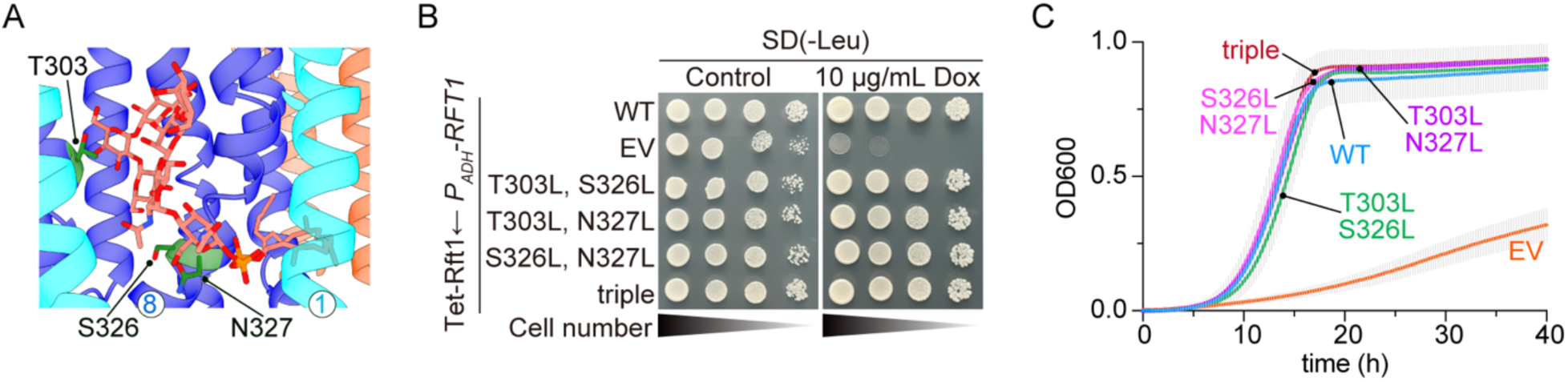
Test of Rft1 with mutations predicted to affect M5-DLO glycan binding. **A.** Rft1-M5-DLO model (Fig. 2A, occluded conformation) showing the positions of residues T303, S326, and N327. TM helices 1 and 8 are indicated as numbers within white circles. **B.** Serial 10-fold dilutions of Tet-off Rft1 cells transformed with an empty vector (EV), wild-type Rft1 (Rft1), or the indicated multi-site point mutants under control of the ADH promoter were spotted onto SD(−Leu) plates with or without 10 μg/mL Dox. The plates were incubated at 30°C for 3 days before being photographed. **C.** Growth at 30°C of the same cells as in panel B, in the presence of Dox, obtained as described in Fig. 4B,C (line and error bars correspond to the mean and standard deviation of 6 measurements). Doubling times for all traces (except EV) range from 190-201 min.

Of the 21 single mutants that we tested, two – R334D and G424Q (Fig. 7A) – did not support normal growth of the Tet-off strain on Dox (Fig. 7B,D). These mutants were designed to modify the central cavity by charge inversion (R334D) and steric occlusion (G424Q) (Fig. 3). Both mutant proteins were similarly well-expressed using the *P_ADH_-RFT1* plasmid, at 50% (R334D) and 80% (G424Q) of the level of wild-type protein (Fig. 7C). However, in spotting assays on Dox plates (Fig. 7B), Rft1(G424Q) supported growth poorly and Rft1(R334D) only slightly surpassed the EV control. These results were confirmed in assays using Dox-containing liquid media (Fig. 7D). However, when the proteins were expressed at a ∼5-fold higher level by using the stronger *P_GPD_* promoter (Fig. 7C) their ability to support growth improved substantially (Fig. 7B,D). Interestingly, despite the discernibly better growth of cells expressing *P_ADH_-RFT1(G424Q)* versus *P_ADH_-RFT1(R334D)* (Fig. 7B,D), *N*-glycosylation of CPY in the two strains remained similarly poor (Fig. S8).

**Figure 7.**
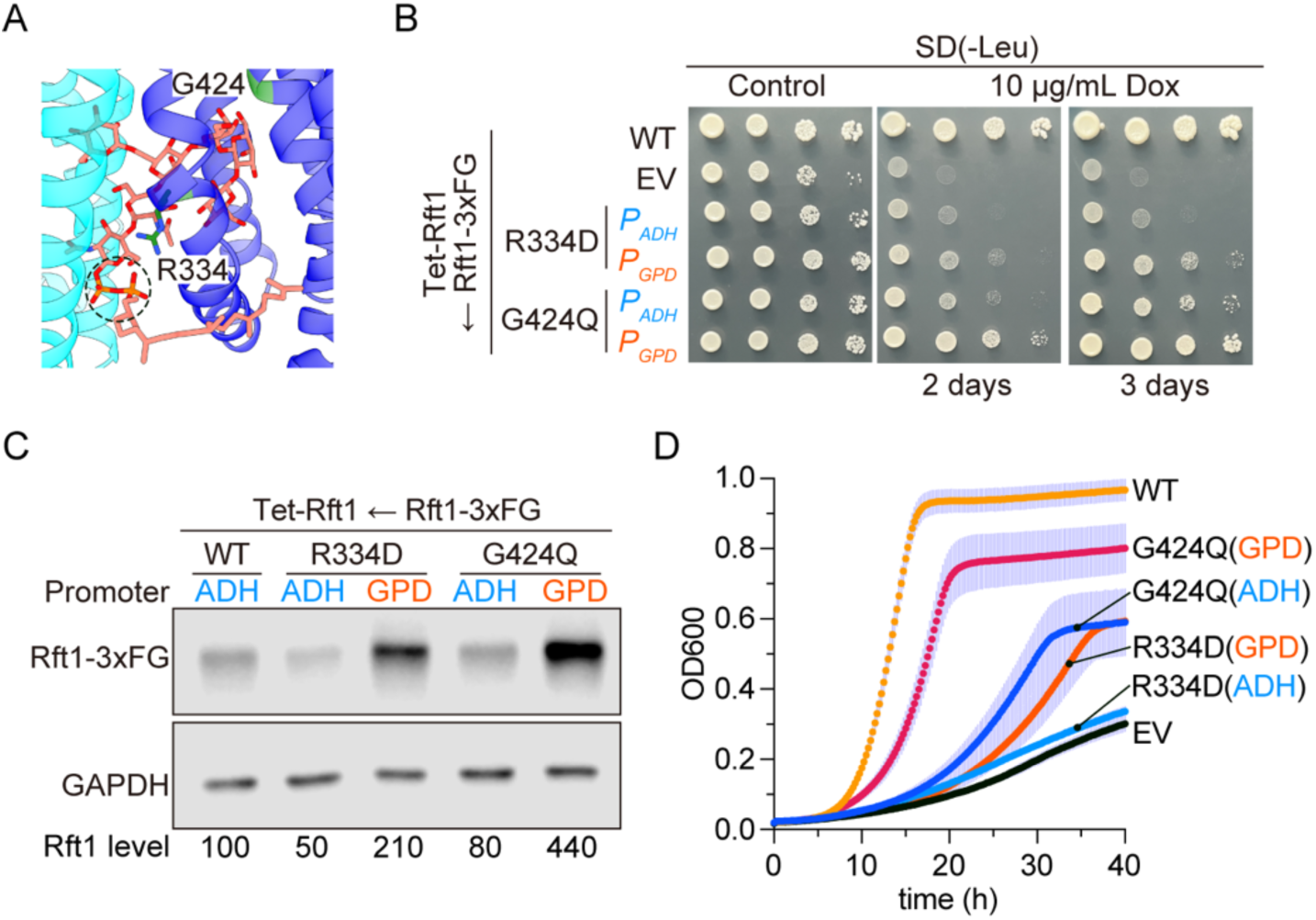
Functional test of R334D and G424Q Rft1. **A.** Rft1-M5-DLO model (Fig. 2A, occluded conformation) showing the proximity of residues R334 and G424 to M5-DLO. **B.** Serial 10-fold dilutions of Tet-off Rft1 cells transformed with an empty vector (EV), wild-type Rft1 (under the ADH promoter), or the R334D and G424Q point mutants (under either the ADH or GPD promoters) were spotted onto SD(−Leu) plates with or without 10 μg/mL Dox. The plates were incubated at 30°C for 2 or 3 days as indicated before being photographed. **C.** Expression levels of Rft1-3xFLAG in the same cells as in panel B (excluding EV). Rft1 level was calculated from band intensities which were normalized to the internal GAPDH control and expressed relative to WT (set to 100%). **D.** Growth of suspension cultures of the same cells as in panel B. Dox (0.5 µg/mL) was added at t = 0 h to the +Dox samples. Data are presented as mean ± SD (n = 6 technical replicates). Doubling times (min) were 180 ± 6 (WT), 1470 ± 190 (EV), 1470 ± 110 (R334D (ADH)), 660 ± 25 (R334D (GPD)), 580 ± 50 (G424Q (ADH)), and 300 ± 9 (G424Q (GPD)). Ordinary one-way ANOVA indicates p<0.0001 for all samples when compared with WT, except for G424Q (GPD) where the difference was not significant.

The results with the R334D and G424Q mutants suggest that over-production of Rft1 variants via the *P_ADH_* and *P_GPD_* promoters may mask defects by mass-action compensation. To address this possibility, we ’under-expressed’ a set of mutants using the *P_RFT1_-RFT1* plasmid (Fig. S9). We found that the K34D, K34W (analogous to human R25W, an RFT1-CDG mutation)^1^, D306K and K307D mutants were able to support growth of the Tet-off cells on Dox albeit to different extents (Fig. S9), whereas R334A (analogous to human R290A, an RFT1-CDG mutation)^1^ grew poorly, similarly to *P_ADH_*-expressed R334D (Fig. 7). Given the already low abundance of Rft1 when expressed from the *P_RFT1_-RFT1* plasmid (∼10 copies per cell for Rft1(WT) on Dox media (Fig. 4D,E)), it is possible that the reduced ability of some of these mutants to support robust growth is due, at least in part, to diminished copy number.

Our data show that the majority of the central cavity mutants robustly support growth of Rft1-deficient yeast cells (Fig. 5), whereas the weak growth-supporting activity of others such as R334D is improved by over-expression. This striking mutational tolerance seemingly contrasts with the situation in MurJ and WzxE, where certain charge-inversion mutations result in total loss of function^29–32,40,41^. It also contrasts with our previous observations of human Rft1 (hRft1)^1^ where the clinical RFT1-CDG mutations R25W and R290A (residues analogous to yeast K34 and R334) failed to support growth despite being well-expressed^1^. It would appear that the central cavity of yeast Rft1 is relatively plastic in contrast with that of the human protein, or that of the bacterial flippases, possibly because it is expansive: in the inward-open state it is >10-times the volume of the M5-DLO headgroup. This volumetric clearance would enable the mutants to physically sequester M5-DLO upon sealing the gating portals, permitting sufficient substrate engagement, e.g., via R334, to execute Rft1’s essential survival function.

### Cell growth is supported by the conformationally locked F38D/Y328K portal mutant

To directly evaluate if M5-DLO scrambling is necessary for Rft1’s ability to support the growth of Tet-off cells on Dox media, a prediction of *hypothesis 1*, we tested the F38D/Y328K double mutant. This mutant is expected to be unable to scramble M5-DLO as it has a hydration-supported salt bridge that prevents the dilation necessary for M5-DLO transit between the fully occluded and outward-facing conformations (Fig. 8A; Fig. S6)^42^. Strikingly, we found that the F38D/Y328K variant successfully supported the growth of the Tet-off cells on Dox plates (Fig. 8B) and in liquid culture (Fig. 8C). Liquid culture assays indicated only a ∼20% longer doubling time for cells relying on the mutant protein compared with those expressing wild-type Rft1 (215 ± 8 min versus 179 ± 4 min; mean ± S.D., n=6) (Fig. 8C). Under these conditions, *N*-glycosylation efficiency remained remarkably high (Fig. S7), with a large fraction of CPY molecules processing the full set of four *N*-glycans. As the double mutant protein expresses as well as the wild-type protein (Fig. S7), we suggest that its slightly depressed activity (growth, *N*-glycosylation) is a result of expected perturbation to the M5-DLO binding site associated with the conformational lock. To stringently test its functionality under restricted abundance, we used the native *P_RFT1_* promoter for expression. Even at the limited level of protein produced under these conditions, the mutant was able to support cell viability, albeit with a four-fold extended doubling time. The remarkable ability of this mutant to sustain cell growth and *N*-glycosylation, despite a mechanically restrictive interhelical tether, strongly implies that M5-DLO scrambling through the lateral portal is dispensable for Rft1’s essential cellular function.

**Figure 8.**
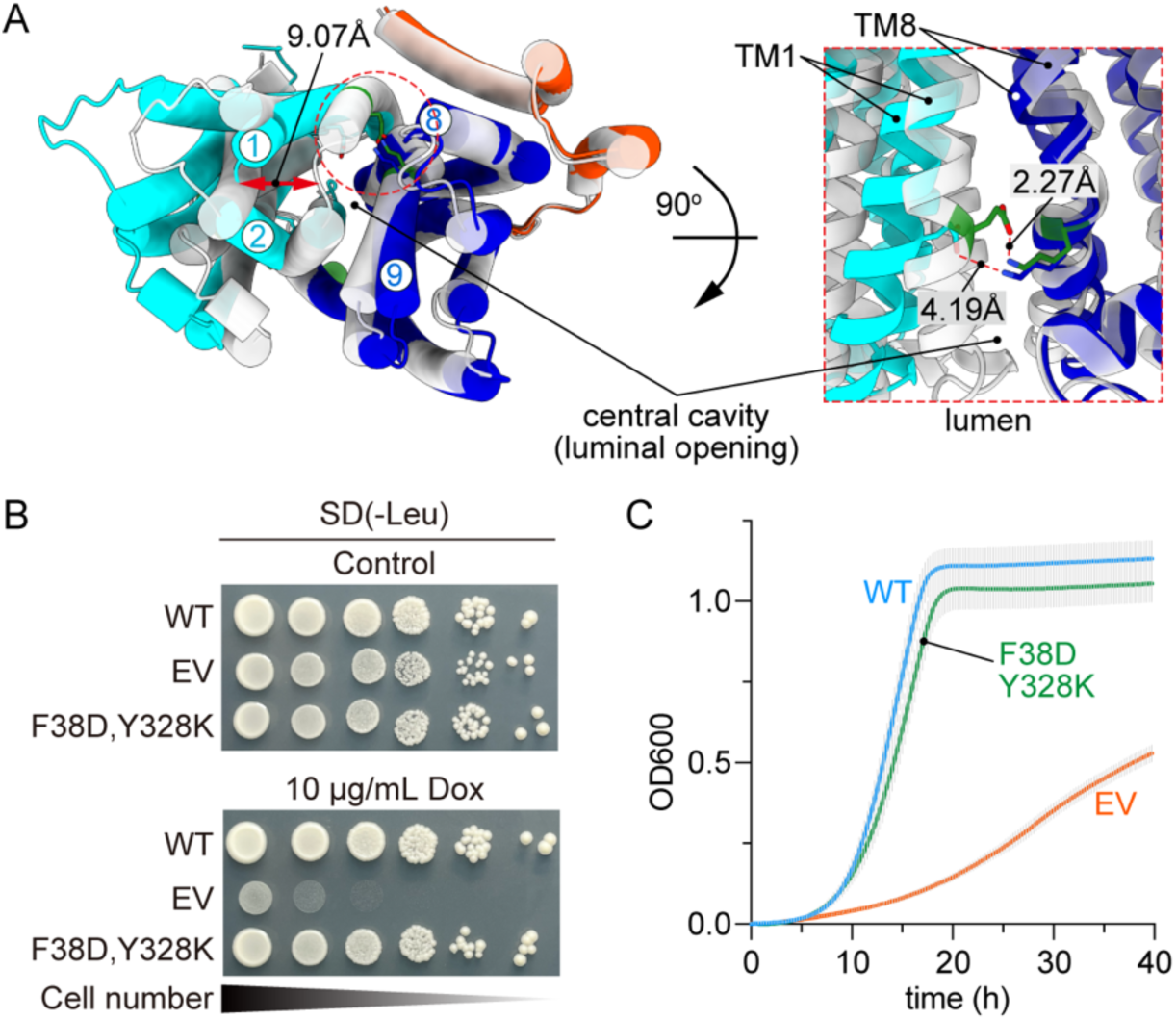
Functional test of the F38D/Y328K ’salt-bridge’ mutant. **A.** Luminal view (left panel) of the outward-facing partially occluded conformation, with the N-lobe in cyan, C-lobe in blue, and additional helices in orange-red, aligned against the occluded conformation in transparent grey. The displacement of TMs 1 and 2 between the occluded and outward-facing partially occluded states (double-headed arrow) is 9.07 Å, measured between the C_α_ of L48 located in the middle of the interconnecting loop. The salt bridge is shown within the dashed circle. The right panel shows an enlarged side view of the engineered salt bridge (portal mutant), highlighting the predicted distance between D38 (OD1) and K328 (NZ). Residues D38 and K328 are shown as sticks, colored green in the occluded conformation and cyan and blue in the outward-open conformation. **B.** Serial 10-fold dilutions of Tet-off Rft1 cells transformed with EV, wild-type Rft1, or the F38D/Y328K mutant were spotted onto SD(−Leu) plates with or without 10 μg/mL Dox. The plates were incubated at 30°C for 4 days before being photographed. **C.** Growth at 30°C of the strains shown in panel B, in the presence of Dox, obtained as described in Fig. 4B,C (line and error bars correspond to the mean and standard deviation of 6 measurements).

## Discussion

Phylogenetic analysis indicates that Rft1 proteins constitute a deep evolutionary lineage distinct from bacterial flippases (Fig. S1). Despite this divergence, AlphaFold3 and Chai-1 modeling reveals that Rft1 retains the core MOP fold found in transporters such as the 12-TM Lipid III flippase WzxE and shares the unique 14-TM architecture characteristic of the MurJ subfamily of Lipid II flippases. Our structural models suggest that Rft1 accommodates the anionic headgroup of its M5-DLO substrate within a voluminous, cationic central cavity, while the isoprenoid tail extends through a lateral portal to associate with a hydrophobic groove on the membrane facing surface of the protein^42,50^ (Fig. 3A). Based on MOP-superfamily architecture, Rft1 would translocate M5-DLO bidirectionally via a MurJ-like alternating-access mechanism (Fig. 2A), distinguishing it energetically and structurally from "credit card" scramblases that use hydrophilic grooves to facilitate continuous lipid sliding^39,42,51,52^. Experimental validation of this mechanism will require state-resolved structural models in conjunction with activity assays, an objective for future work.

We present two hypotheses to account for Rft1’s essentiality in yeast and human cells, both of which require M5-DLO to bind to Rft1. For *hypothesis 1*, binding is a prelude to essential scrambling via the alternating access mechanism, whereas for *hypothesis 2* it is an element of the ’catch and release’ activity that underlies Rft1’s proposed role as an M5-DLO chaperone. Scrambling is not precluded by *hypothesis 2*, but it is a moonlighting function secondary to the essential chaperone function, and redundant with the activity of other cellular scramblases.

Our results align with *hypothesis 2*. The poor growth-supporting activity of the R334D and G424Q variants (Fig. 7) indicates that M5-DLO binding via the central cavity is indeed important for Rft1’s function as posited. This point is reinforced by the inability of analogous RFT1-CDG mutants (R25W, R290A) to support growth of yeast cells despite being expressed at levels comparable to that of the wild-type hRft1 protein. The key result that the F38D/Y328K portal mutant supports cell growth argues strongly against *hypothesis 1*. Engineered to introduce a hydration-supported interhelical salt bridge, this variant has a rigid conformational lock to obstruct the lateral portal and stall the transport cycle (Fig.7, Fig. S6). Remarkably, even under restricted expression using the *P_RFT1_* promoter, the F38D/Y328K variant sustained cell viability indicating that transit of M5-DLO between the occluded and outward-facing states is not required for Rft1’s function in cells.

Considerations of transport kinetics provide further support for *hypothesis 2*. Glycoproteomics data^53^ indicate that a yeast cell has ∼500 *N*-glycoproteins. Based on the number of observed glycopeptides^53^ and protein copy number estimates^49^, this corresponds to ∼14 million *N*-glycans per cell. With a typical doubling time for yeast cells being 90 min (the doubling times that we report here are longer because our growth assays were done in multi-well plates), we calculate that 155,000 *N*-glycans must be synthesized per minute and thus infer that 2,600 M5-DLO molecules must be generated and scrambled per second to cope with glycosylation requirements. As there are ∼20 copies of Rft1 per Tet-off cell, Rft1’s turnover rate must be 130 M5-DLO molecules per second if it is the only M5-DLO scramblase in cells. The alternating access mechanism of MOP family proteins involves large-scale, intra-membrane, rigid-body helical rearrangements which inherently restrict turnover rates^29,54,55^; for example, the estimated turnover rate for MurJ is only ∼10 Lipid II molecules per second^56^. However, if Rft1 functions as a cytoplasmic chaperone, rapid substrate on/off binding kinetics (uncoupled from the slow transmembrane inversion) may easily allow the ∼20 copies of Rft1 to process the required lipid flux. These back-of-the-envelope calculations suggest that Rft1 is unlikely to meet the demand for M5-DLO flipping in cells and that additional scramblases are necessary^6,7^.

In summary, the confluence of our structural, genetic, and kinetic analyses challenges the hypothesis that Rft1’s essential function in cells is to scramble M5-DLO. Instead, we propose that Rft1’s essential function is that of a spatial chaperone, a role in which its central cavity is strictly required to capture M5-DLO. Thus, Rft1 would ferry M5-DLO within the plane of the ER membrane or precisely hand it off to downstream biosynthetic machinery or redundant scramblases, thereby ensuring cellular viability.

## Methods and methods

### Rft1 structure-based phylogeny

The AlphaFold model (AF-P38206-F1-v6) was downloaded from the AlphaFold database via the Uniprot entry (P38206) for *Saccharomyces cerevisiae* Rft1^57,58^. Sequence conservation analysis was performed using the default parameters and mapped onto the AlphaFold model through the ConSurf server using default parameters^59–62^. Rft1 structural homologues were identified by searching the Protein Data Bank with a Clustal O^63^ multiple sequence alignment of Rft1 sequences from *Saccharomyces cerevisiae, Trypanosoma brucei*, *Homo Sapiens*, *Drosophila Melanogaster*, *Danio rerio* and *Caenorhabditis elegans* using the HHpred server^33^. For structure-based phylogenetic reconstruction a dataset of 27 representative AlphaFold Database protein structure predictions comprising Rft1 orthologs (fungal, metazoan, and protozoan) and bacterial lipid flippases was compiled based on the results of independent FoldSeek^64^ search results queried using HHpred hits *Escherichia coli* WxzE and *Thermosipho africanus* Murj as well the yeast Rft1. Due to the low sequence identity between eukaryotic and bacterial MOP family members (typically <15%), standard sequence-based alignment methods are prone to alignment errors in the "twilight zone" of homology. Therefore, we utilized FoldMason^65^, a structure-based multiple sequence alignment tool that leverages the 3Di structural alphabet to generate high-fidelity alignments based on tertiary structure conservation rather than primary sequence alone.

The resulting structure-guided amino acid alignment (717 columns) was used for phylogenetic inference using IQ-TREE (version 1.6.12)^66^. Model selection was performed using ModelFinder^67^, which identified the LG+F+G4 model (Le and Gascuel matrix with empirical base frequencies and 4-category Gamma rate heterogeneity) as the best-fit substitution model based on the Bayesian Information Criterion (BIC). While compositional heterogeneity was observed in 30.3% of the sequences (Chi-square test, p < 0.05), alternative profile mixture models (e.g., C20) resulted in significantly poorer fit scores compared to the standard LG model in preliminary runs, indicating that the LG+F+G4 model provided the most robust fit for this dataset. Branch support was assessed using 1000 replicates of both the SH-like approximate likelihood ratio test (SH-aLRT)^68^ and Ultrafast Bootstrap (UFBoot)^69^. Tree visualization was done using iTOL (v7.5.1)^70^.

### M5-DLO docking and ligand topology

The structure prediction algorithms Alphafold 3^26^ (installed locally) and Chai-1^27^ (accessed through the web portal) were used to predict alternate conformations of Rft1 and the Rft1/M5-DLO complex. The initial structure of M5-DLO was download from the Chemical Entities of Biological Interest data base, entry CHEBI:132515 and converted to SMILES format^71^. Instead of the typical 14-19 isoprenyl repeats for yeast M5-DLO^72^, a shortened version with four of these was used to simplify the docking analysis. Our choice of a short C_20_ isoprenoid to model M5-DLO is supported by recent data showing that Rft1 can scramble Man_5_GlcNAc_2_-PP-phytanol which contains a C_20_ chain^15^. Chai-1 was run without multiple sequence alignment and no constraints, producing (by default) five Rft1/M5-DLO complex models. Residues within 5 Å of the docked M5-DLO in each of the five Rft1/M5-DLO models were determined using the Residue Interaction Network analyser and structureViz plugins in Cytoscape (v 3.10.3) and Chimera (v1.19)^73–76^. Five seeds were used for the local AlphaFold3 run, with M5-DLO specified in SMILES format, generating 25 models of Rft1/M5-DLO complexes.

To enforce stereochemical accuracy, particularly at the critical lipid-diphosphate-glycan interface, a second AlphaFold3 run was performed where ligand topology was defined using the Chemical Component Dictionary (CCD) format linked via the bondedAtomPairs syntax, rather than SMILES strings, which have been shown to frequently misassign anomeric configurations in complex glycans^36^. The M5-DLO ligand was assembled using octaprenyl pyrophosphate (CCD code: OTP) as a structural analog for the dolichol carrier. The glycan moiety was constructed from standardized monosaccharide building blocks. To recapitulate the native biosynthetic linkage, *N*-acetyl-α-D-glucosamine (CCD code: NDG) was explicitly specified as the reducing-end residue connected to the pyrophosphate, preserving the biological α-anomeric configuration distinct from protein *N*-glycosylation. This was extended by *N*-acetyl-β-D-glucosamine and β-D-mannose to form the core, with subsequent β-D-mannose branches defined via explicit atom-pair bonding to match the canonical M5 topology. Modeling was performed using three independent random seeds. The correct stereochemistry of the docked M5-DLO in all the 15 complex models were assessed and confirmed to be realistic using the Privateer online tool^77,78^. Chimera X (v1.10) was used for structure visualization, alignment and figure preparation.

### Volumetric analysis of the central cavity

To evaluate the macroscopic dimensions of the putative central M5-DLO binding cavity, volumetric analyses were conducted across the computationally predicted states of the Rft1 transport cycle. Measurements were performed using the CASTpFold web server, which analytically computes solvent-accessible surface area and internal volume. The algorithm was executed using default parameters, applying a standard 1.4 Å water probe to precisely define the accessible boundaries of the internal cavity^37^. MoloVol was used to calculate the size of the polar head group of M5-DLO as modeled by AF3 using CCD/BPA syntax^38^ .

### Molecular biology

Rft1 point mutants driven by the ADH promoter were generated by inverse PCR using p415[ADHpro-yRft1-3×FLAG] (EcAKM406, *LEU2* marker) as a template. For R334D and G424Q, additional constructs were made in which the ADH promoter was replaced with the GPD promoter using restriction enzyme-based cloning. Rft1 point mutants driven by the native Rft1 promoter were constructed by amplifying the *RFT1* ORF fragments from the ADH promoter–driven mutant plasmids by PCR, followed by replacement of the *RFT1* ORF in pRS315[Rft1–3×FLAG] (EcAKM465, *LEU2* marker) using Gibson assembly. All mutations were verified by Sanger sequencing. To construct pRS315[Rft1–3×FLAG], a DNA fragment containing the *RFT1* ORF along with 500 bp upstream and 450 bp downstream sequences was amplified from BY4741 genomic DNA and inserted into the SmaI site of the pRS315 vector to generate pRS315[Rft1] (EcAKM222). A sequence encoding a 3×FLAG tag was subsequently inserted immediately upstream of the stop codon of the *RFT1* ORF.

### RT-qPCR

Tet-Rft1 cells carrying an empty vector (EV), or vectors encoding Rft1-3xFLAG (expression driven by *P_RFT1_* or *P_ADH_* promoters, were cultured to mid-log phase (OD600=0.8), and 10 OD600 units were collected for mRNA isolation. The cells were washed with 1.2 M sorbitol (Sigma S7547), 50 mM Na_2_HPO_4_, 20 mM EDTA, 1% β-mercaptoethanol, resuspended in 2 ml of same buffer containing 25 µg of zymolyase 20T (amsbio 120491-1) and incubated for 30 min at 30°C. Cells were pelleted by centrifugation (300 × g, 5 min) and resuspended in 1 mL TRIzol reagent (Invitrogen 15596026). Samples were stored at −80°C overnight prior to RNA extraction.

Total RNA was isolated using a modified TRIzol protocol. Briefly, samples were vortexed at room temperature for 5 min, followed by addition of chloroform and separation of phases by centrifugation (15,000 rpm, 5 min, 4°C). The aqueous phase was transferred to a new tube and subjected to a second chloroform extraction. RNA was precipitated by addition of 0.5 volumes of isopropanol and incubation at -20°C for 15 min, followed by centrifugation (15,000 rpm, 5 min, 4°C). The RNA pellet was washed twice with ice-cold 70% ethanol, briefly air-dried, resuspended in 50 µL DEPC-treated water, and aliquoted and stored at -80°C until use.

Aliquots of total RNA were used for RT–qPCR analysis. Genomic DNA was removed using gDNA Eraser (Takara, RR047A), and cDNA was synthesized using the PrimeScript RT Reagent Kit (Takara, RR047A) according to the manufacturer’s instructions. Quantitative PCR was performed using SYBR Green Universal Master Mix (Applied Biosystems, 4309155) under the following conditions: initial denaturation at 95°C for 20 s, followed by 40 cycles of 95°C for 3 s and 60°C for 30 s. Amplification was monitored using a C1000 Touch Thermal Cycler and a CFX96 Touch Real-Time PCR Detection System (Bio-Rad).

All experiments were performed with three independent biological replicates, and each sample was analyzed in technical duplicates. Relative gene expression levels were calculated using the ΔΔCt method with PGK1 as an internal control. Ct values of target genes were normalized to PGK1 to obtain ΔCt values. ΔΔCt values were calculated relative to the EV control, and fold changes in gene expression were determined as 2^−ΔΔCt^.

Primer sequences used for qPCR were as follows: RFT1, forward 5ʹ-AGCGAGCCATTCTTCATCGT-3ʹ, reverse 5ʹ-TAGTCACCGCGATGCTTTCA-3ʹ; PGK1, forward 5ʹ-TCTTCGACAAGGCTGGTGCTGAAA-3ʹ, reverse 5ʹ-ACAAAGCCTTAGTACCAGCAGCGA-3ʹ.

### Yeast growth assays

Plasmids were introduced into the Tet-off-RFT1 strain (YAKM147: *URA3::CMV-tTA, his3-1 leu2-0 met15-0 RFT1::kanR-TetO_7_-CYC1_TATA_-RFT1*). Transformants were selected on an SD(-Leu) plate. For growth assays on plates (spotting assays), colonies were resuspended in sterile water and adjusted to OD600 = 1.0. Ten-fold serial dilutions were prepared and 10 µL of each dilution was spotted onto SD(-Leu) plates with or without 10 µg/mL Dox. Plates were incubated at 30°C for 2–3 days before being photographed. For analysis of the growth of cells in suspension, growth curves were generated essentially as described previously^1^. Briefly, transformed cells were pre-cultured overnight in SD(-Leu) medium, diluted into fresh SD(-Leu) medium, and grown to mid-log phase (OD600 = 0.4–0.8). Cells were then diluted into YPD + Dox (0.5 µg/mL), OD600 = 0.01, and dispensed into a 96-well plate (200 µL per well). The plate was sealed with a Breathe-Easy polyurethane membrane (Sigma-Aldrich, Z380059) and incubated at 30°C in a SpectraMax i3x plate reader (Molecular Devices). OD600 was recorded every 15 min for 40 h with 5 s of orbital shaking prior to each measurement. Doubling time was calculated as previously described^1^, where C₁ and C₂ were chosen within the linear portion of the growth curve, usually OD600 = 0.5–0.65.

### Protein extraction and immunoblotting

This was done as described previously^1^. Briefly, cells were cultured to mid-log phase, and 1 OD600 unit of cells was collected. Cells were lysed in 0.28 mL of 0.2 M NaOH containing 0.5% β-mercaptoethanol and incubated on ice for 5 min. Proteins were precipitated by adding 0.5 mL of 30% trichloroacetic acid (TCA) and incubating on ice for 10 min. Precipitated proteins were collected by centrifugation (14,000 rpm, 5 min, 4°C), washed twice with 0.2 mL of ice-cold acetone, air-dried, and solubilized overnight in 50 µL of 2% SDS, 5 mM NaOH, and 2% β-mercaptoethanol. The samples were mixed with 50 µL of 2× SDS-PAGE loading buffer and subjected to SDS-PAGE/immunoblotting. CPY, GAPDH, and Rft1-3xFLAG variants were detected with anti-CPY (1:5000; Invitrogen 10A5B5), anti-GAPDH (1:100,000; Sigma G9545), and anti-FLAG (1:5000; Sigma-Aldrich F1804) antibodies and visualized using horseradish peroxidase-conjugated secondary antibodies anti-mouse IgG (1:10,000; Promega W402B) and anti-rabbit IgG (1:10,000; Promega W401B). Signals were visualized using an Odyssey XF imaging system (LI-COR), and band intensities were quantified with Image Studio software (LI-COR). For assessing the expression level of Rft1-3xFLAG constructs, the FLAG signal was normalized to that of GAPDH. The copy number of Rft1-3xFLAG per cell was determined by quantifying the FLAG immunoblot signal with that of a series of 3xFLAG-tagged bovine opsin standards of known concentration. Glycoscores were calculated as described previously^1^ to provide a measure of CPY *N*-glycosylation.

## Author contributions

George N. Chiduza: conceptualization, data curation, formal analysis, investigation, methodology, validation, visualization, writing – original draft preparation, writing – review and editing; Kentaro Sakata: investigation, formal analysis, visualization, writing – review and editing; Faria Noor: investigation; Hannah G. Wolfe: investigation; Anant K. Menon: conceptualization, data curation, formal analysis, funding acquisition, supervision, validation, visualization, writing – original draft preparation, writing – review and editing.

## Supplementary material description

Supplementary figures (Figs. S1-S9) and tables (Tables S1-S2) accompany this article.

The results of computational analyses are uploaded to Mendeley Data, doi: 10.17632/ns9n5mjy8x.1

## Acknowledgement

We acknowledge support from National Institutes of Health grant R01 GM146011 (A.K.M.), and the Fonds National de la Recherche Scientifique (FRS-FNRS; Grant number 1.B.089.24F) (G.N.C.). We thank Maya Schuldiner (Weizmann Institute of Science) for the Tet-off RFT1 strain, Indu Menon (Weill Cornell Medicine) for a sample of 3xFLAG-tagged bovine opsin, Ronghu Wu (Georgia Institute of Technology) for pointing us to yeast glycoproteomics data, Baran Ersoy (Weill Cornell Medicine) for use of a plate reader, Ariel Talavera Perez (Université Libre de Bruxelles) for managing computational resources, Rie Nygaard and Olga Boudker (Weill Cornell Medicine), Jean-Marie Ruysschaert (Université Libre de Bruxelles), and Jeff Rush and Skip Waechter (University of Kentucky) for comments on the manuscript,.

