## Supplementary Information for "Mechanism-guided mutagenesis of Rft1 to test its role as a dolichol-linked oligosaccharide scramblase in cells"

Supplementary Information
Figures S1-S9
Tables S1, S2

### Mechanism-guided mutagenesis of Rft1 to test its role as a dolichol-linked oligosaccharide scramblase in cells

George N. Chiduza[1,*], Kentaro Sakata[2], Hannah G. Wolfe[2], Faria Noor[2] and Anant K. Menon[2]

[1]Biochemistry and Structural Biology-Chemistry Department, Université Libre de Bruxelles - Campus Plaine, 1050 Brussels, Belgium
[2]Department of Biochemistry and Biophysics, Weill Cornell Medical College, New York, NY 10065, USA

*Correspondence
Dr. George N. Chiduza
Biochemistry and Structural Biology- Chemistry Department, Université Libre de Bruxelles- Campus Plaine, 1050 Brussels, Belgium

**Figure S1. Structure-guided maximum likelihood phylogeny of the MOP transporter superfamily.** The phylogenetic tree displays the evolutionary relationships among 27 representative sequences comparing eukaryotic Rft1 orthologs with characterized bacterial MOP superfamily members including MurJ, WzxC/E, and NorM. Eukaryotic Rft1 sequences cluster into a distinct clade (100% UFBoot/SH-aLRT). Within the Rft1 lineage the internal branching depicts the established organismal phylogeny. The bacterial transporters are grouped into a separate superclade partitioned into distinct subgroups, including the MurJ lipid II flippases and the WzxC/E lipid III and O-antigen flippases. The phylogeny was reconstructed in IQ-TREE using a FoldMason structure-guided sequence alignment (717 columns, 589 parsimony-informative sites) to compensate for low overall sequence identity among the distant homologs. The LG+F+G4 substitution model was applied, with branch support calculated from 1000 replicates for both Ultrafast Bootstrap and SH-aLRT. The scale bar indicates amino acid substitutions per site. Tree visualization was done using iTOL 7.5.1 (see Methods).

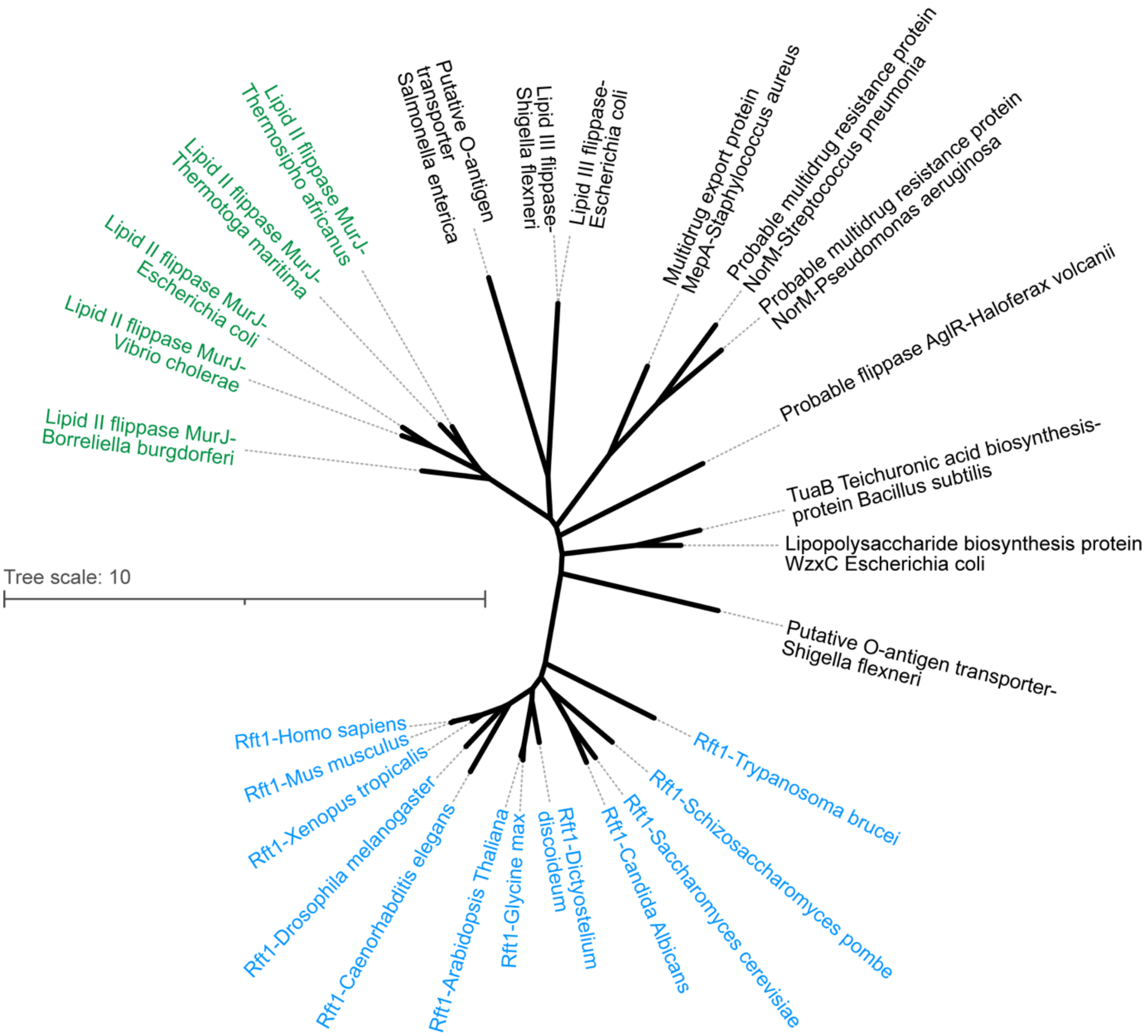

**Figure S2. Comparison of Rft1's predicted structure with MurJ. A.** Model of Rft1 downloaded from the Alphafold data base (AF-P38206-F1-v6), colored by predicted local-distance difference test (pLDDT) confidence score. **B.** Structural alignment of the Alphafold Rft1 model with the crystal structure of *Thermosipho africanus* MurJ in the inward-open conformation (PDB ID: 6NC7). **C.** Chai-1 model of TA-MurJ-Lipid II complex (Le4; Cα-RMSD = 0.983 Å to 6NC7). Lipid II modelled using SMILES from PubChem CID 46173749, shown as green sticks. Charge inverting mutations in R18, R25, R52 and R255 are lethal as they lead to total loss of function in MurJ (right).

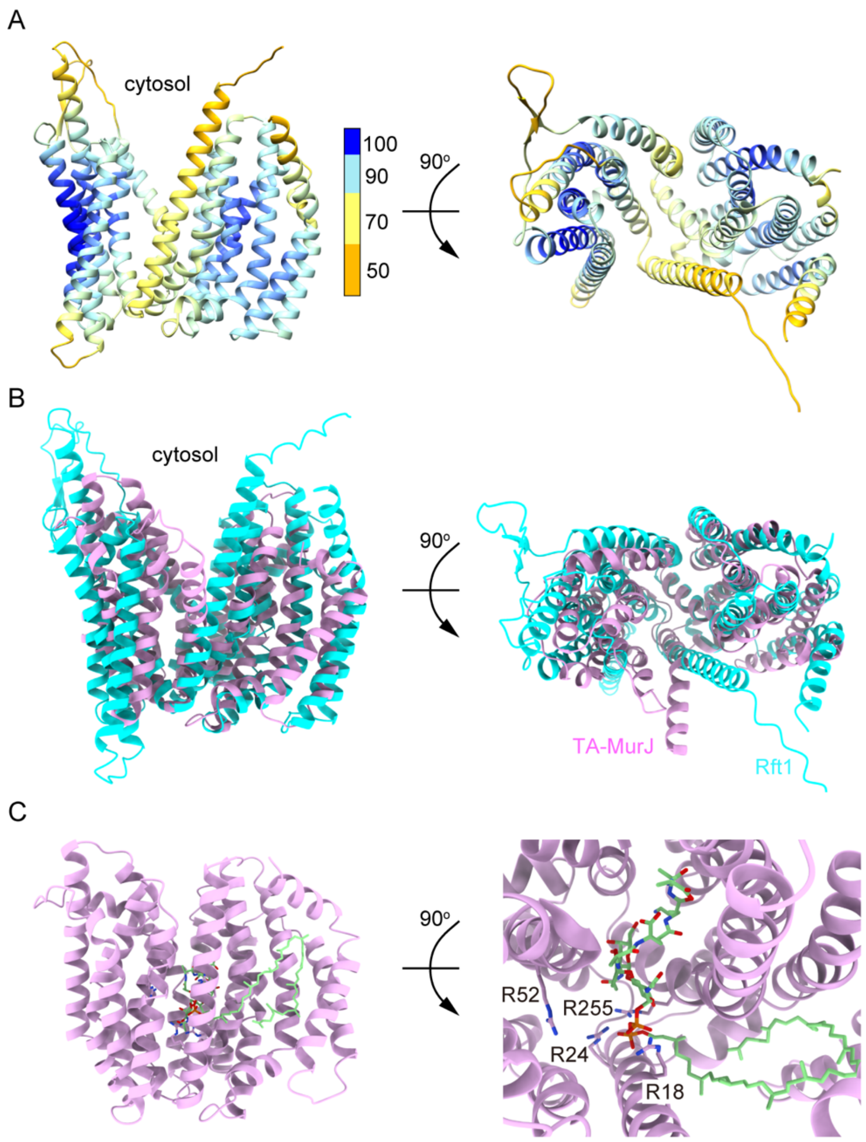

**Figure S3. Sequence alignment used for HHpred search.** Clustal O multiple sequence alignment of Rft1 sequences from *Saccharomyces cerevisiae*, *Trypanosoma brucei*, *Homo Sapiens*, *Drosophila Melanogaster*, *Danio rerio* and *Caenorhabditis elegans*.

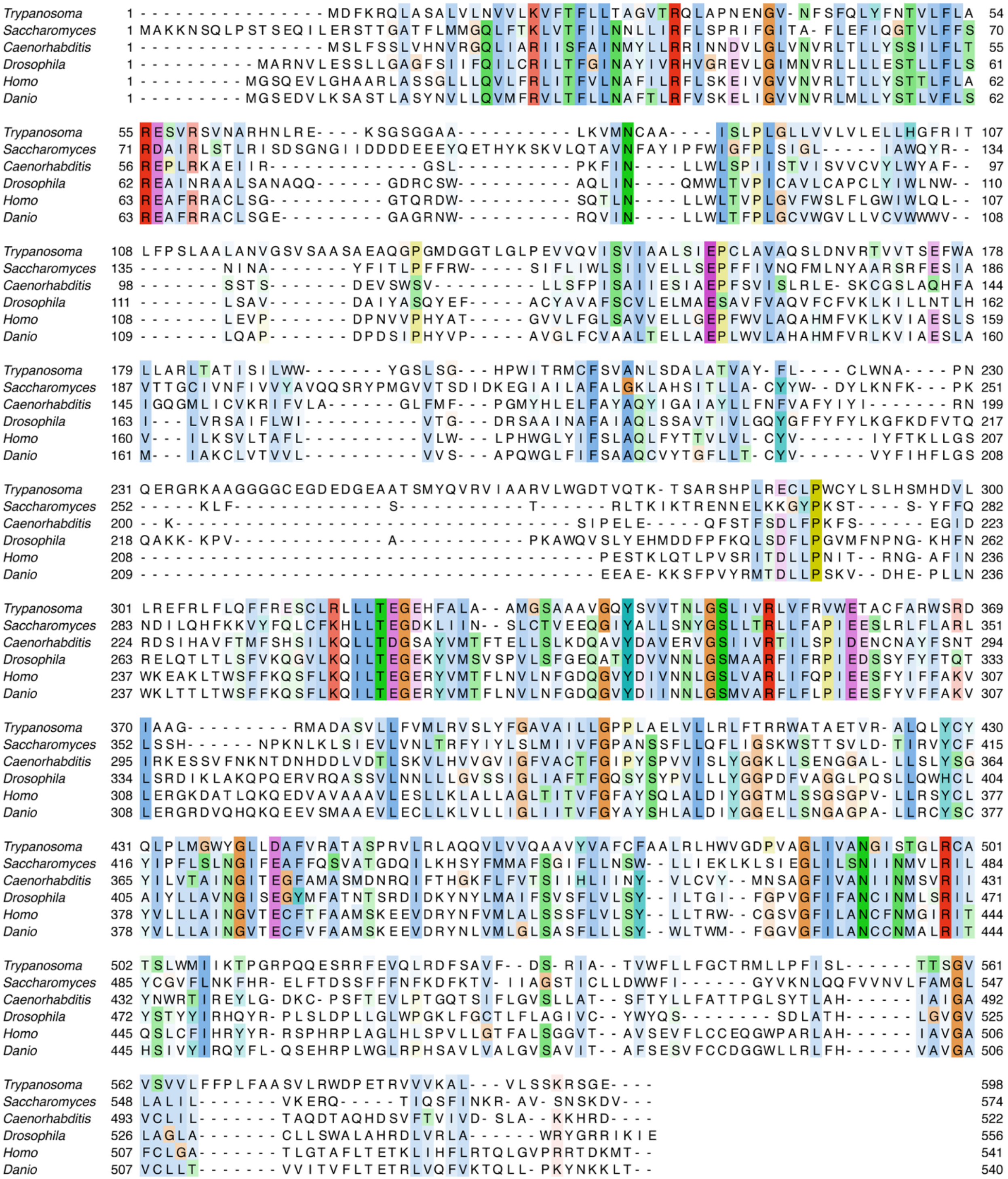

**Figure S4. Stereochemistry of the M5-DLO model generated by co-folding protein structure predictors.**
**A.** M5-DLO defined by Alphafold3 using a CCD/BPA hybrid syntax. **B, C.** Models of M5-DLO produced by Alphafold3 (B) and Chai-1 (C), using SMILES as inputs for the ligand description.

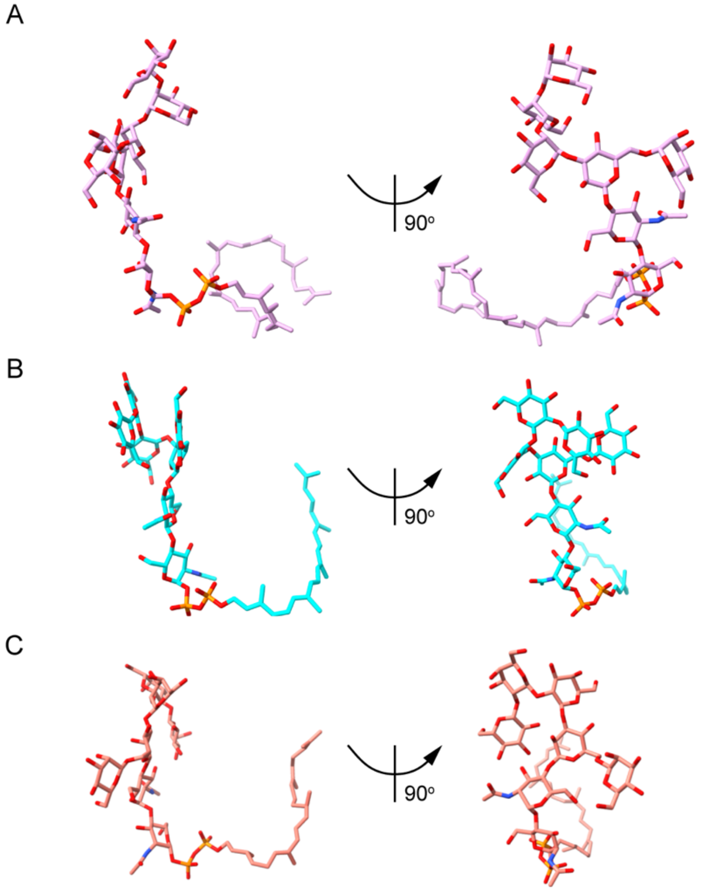

**Figure S5. Volumetric analysis of the Rft1 central cavity across conformational states predicted by Chai-1.** The dimensions of the central binding cavity were calculated using CASTp with a standard 1.4 Å probe radius. **A.** The predicted inward-open (cytosol-facing) state, with a central cavity volume of 12,789 Å³. **B.** The occluded state, displaying a reduced central cavity volume of 6,178.4 Å³. **C.** The lumen-facing state, with a central cavity volume of 8,963.2 Å³.

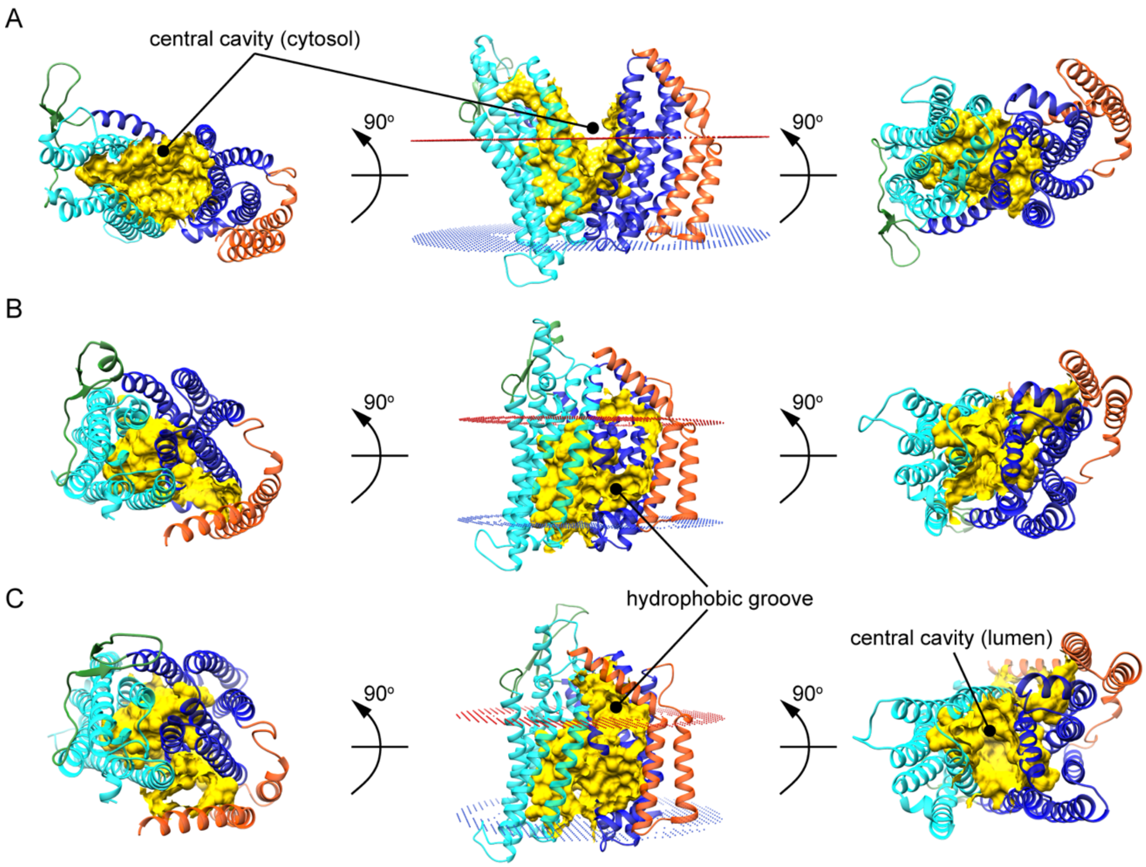

**Figure S6. Schematic illustration of the effect of the F38D-Y328K portal salt bridge on M5-DLO scrambling.** The illustration is labeled according to the inward-open (apo) (I), inward-open (bound) (II), occluded (III) and outward-facing (IV) conformations depicted in Figure 2. Color coding is the same as that shown in Figure 2, with the N-lobe, C-lobe and TM helices 13 and 14 colored cyan, blue and red, respectively. Residues D38 in TM1 and K328 in TM8 are indicated by green circles, connected by a salt bridge (dashed line). M5-DLO entering on the cytoplasmic side (top left) cannot be discharged on the luminal side of the membrane as shown. Luminal M5-DLO (bottom right) can interact with conformation IV but cannot enter the central cavity. The salt bridge restricts Rft1's dynamics to the cytoplasmic leaflet, where it remains fully capable of catching and releasing M5-DLO

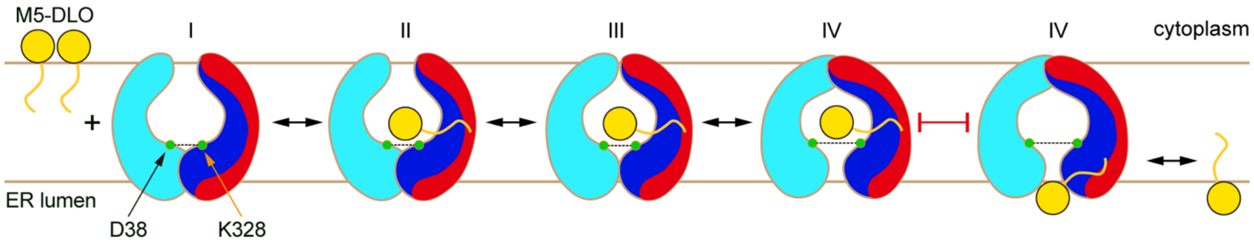

**Figure S7. Expression of Rft1 mutants.**

**A.** Anti-FLAG (to detect Rft1-3xFLAG variants) and anti-GAPDH (loading control) immunoblots. Cells (expressing Rft1 constructs via a $P_{ADH}$-*RFT1* plasmid with a *LEU* marker, were grown on SD(-Leu) media to log phase before collecting samples for immunoblotting analysis. **B.** Expression levels were assessed by quantifying band intensities, taking the ratio of the FLAG signal to the GAPDH signal in each case, and further normalizing to the corresponding ratio for a wild-type Rft1-3xFLAG (set to 100%) analyzed alongside. The bar charts show the mean value of several measurements. The indicated data points correspond – in most cases – to at least 2 biological replicates, with additional points corresponding to technical replicates, and the vertical lines represent the range of the measurements.

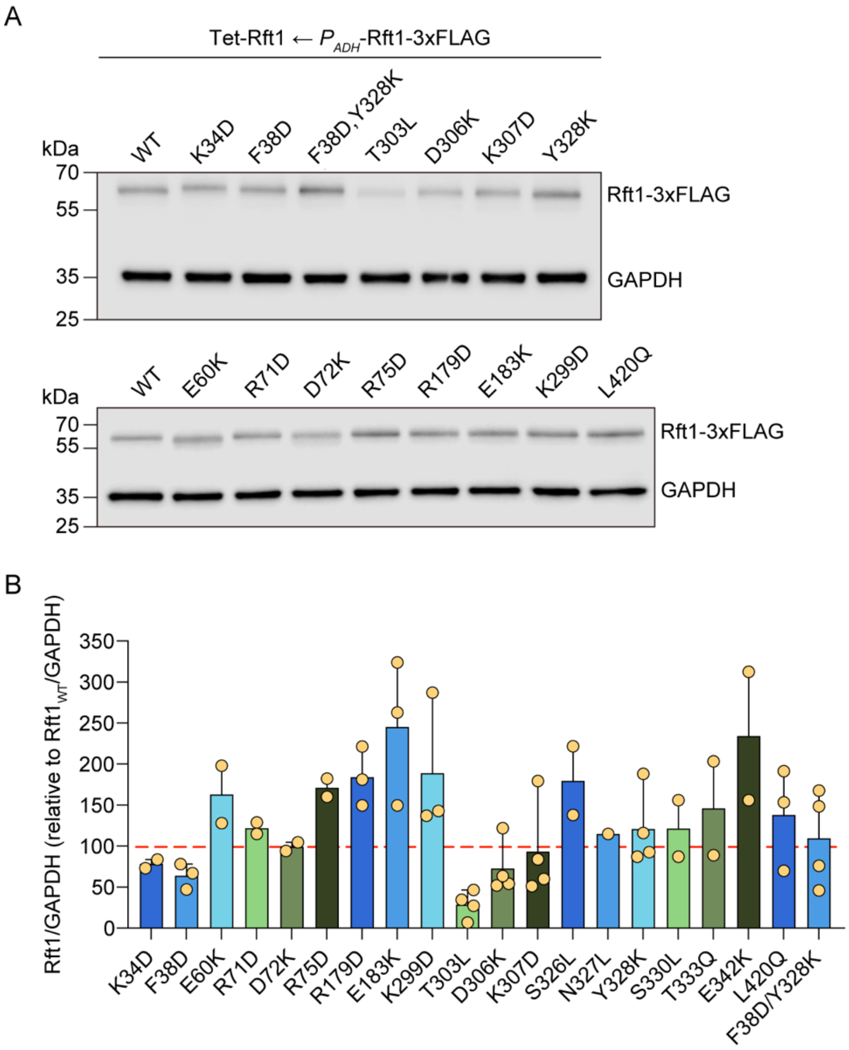

**Figure S8. *N*-glycosylation in cells expressing Rft1 variants.** Tet-off cells were transformed with *LEU* plasmids for expression of Rft1 variants under control of the $P_{ADH}$ or $P_{GPD}$ promoters (as indicated) and grown in SD(-Leu) media before being shifted to YPD media with Dox (0.5 μg/mL) for 17 h at 30°C. Cells were harvested and processed for immunoblotting with anti-CPY antibodies. **A.** Representative immunoblots, with WT and EV controls. Images for T303L and L420Q were taken from a separate blot which also included WT and EV controls (not shown). The column of filled circles (gradient of grey) shown in the lower panelalongside the G424Q sample indicates CPY glycoforms with 4. 3, 2, 1 and 0 *N*-glycan(s). **B.** Glycoscores calculated from multiple blots, including the ones shown in panel A. Data represent the mean and standard deviation (n=2-7 technical replicates, including protein preparations from unique cell pellets), with red dashed lines corresponding to the means for the WT and EV samples. Pairwise comparison of all samples to the WT sample was done with ordinary one-way ANOVA. ns, not significant, **p=0.0034, ***p=0.0001, ****p<0.0001.

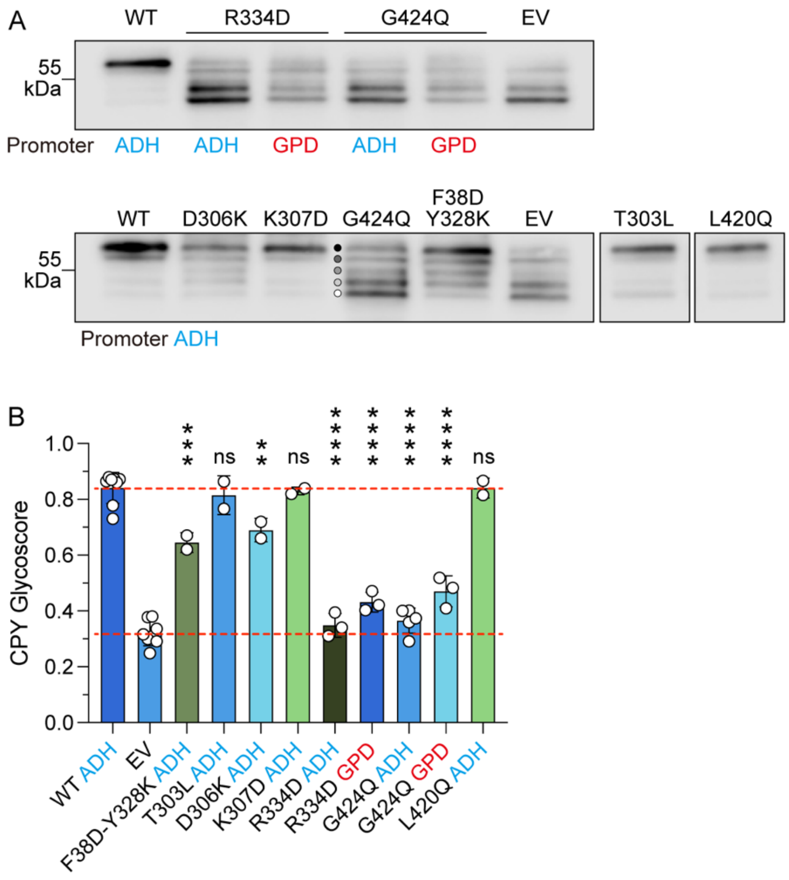

**Figure S9. Testing Rft1 variants expressed under control of the RFT1 promoter. A.** Tet-off cells carrying Rft1 point mutants expressed under control of the $P_{RFT1}$ promoter were serially diluted and spotted onto SD(−Leu) plates with or without 10 µg/mL Dox. The plates were incubated at 30°C for 5 days before being photographed. **B.** Growth in liquid culture at 30°C of the same cells as in panel A, in the presence of Dox, obtained as described in Fig. 4B,C (the lines are the mean of 5-6 measurements). **C.** Doubling times, calculated from the growth curves in panel B. Ordinary one-way ANOVA indicates p<0.0001 for all variants when compared with WT, except for K34D (p=0.0220) and D72K where the outcome was not significantly different.

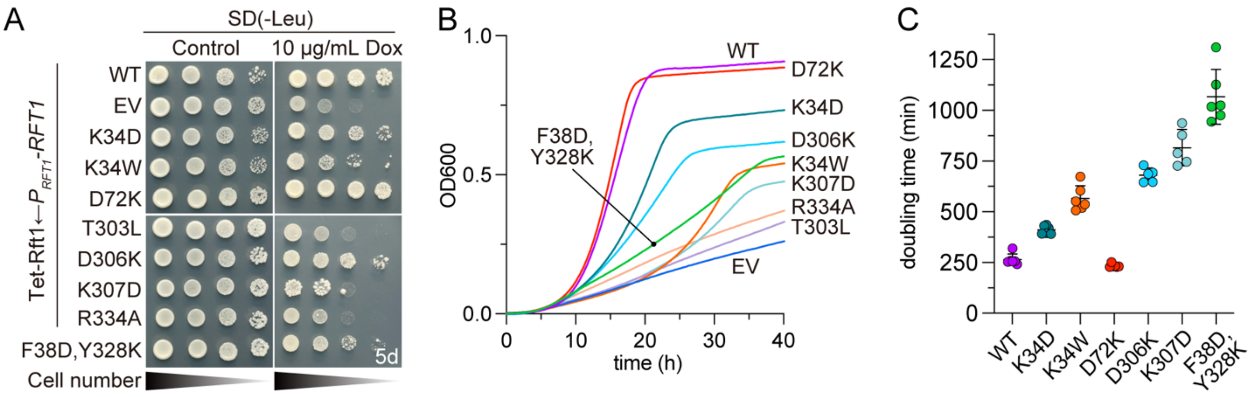

Table S1. Homologues of Rft1 identified by HHPred

| Rank | PDB ID | Name/Description | Probability | E-value | Score | Secondary Structure | Aligned residues | Target Length |
|---|---|---|---|---|---|---|---|---|
| 1 | 7WAX_A | Lipid II flippase MurJ; 14 transmembrane helices, inner membrane, Lipid II, MOP superfamily, LIPID TRANSPORT; HET: OLC, | 99.81 | 9.20E-15 | 137.05 | 42 | 424 | 511 |
| 2 | 9G97_A | Lipid III flippase; membrane protein, flippase, cell wall, enterobacterial common antigen, lipid III, TRANSPORT PROTEIN; | 99.79 | 2.60E-14 | 134.55 | 42.4 | 415 | 425 |
| 3 | 6FHZ_A | Putative MOP flippase; MOP flippase, MEMBRANE PROTEIN; 2.8A {Pyrococcus furiosus DSM 3638} | 99.78 | 1.90E-14 | 132.35 | 38.1 | 428 | 440 |
| 4 | 3W4T_A | Putative uncharacterized protein; MATE, multidrug transporter, TRANSPORT PROTEIN; HET: OLC; 2.096A {Pyrococcus furiosus} | 99.76 | 7.50E-14 | 129.28 | 38.4 | 434 | 461 |
| 5 | 5C6O_A | BH2163 protein; protein binding, TRANSPORT PROTEIN; HET: 4YH; 3.0A {Bacillus halodurans (strain ATCC BAA-125 / DSM 18197 | 99.73 | 4.60E-13 | 124.21 | 39.3 | 431 | 464 |

Table S2

Table S2. Amino acids within 5 Å of M5-DLO in Chai-1 models

| Amino acid position | Consurf Grade | Count* | Amino acid position in human Rft1^ | Phenotype of tested human Rft1 mutant^ | Mutated in RFT1-CDG? |
|---|---|---|---|---|---|
| Q30 | 9 | 3 | Q21L | Glycoscore ~60% | no |
| K34 | 9 | 3 | R25W | Loss of function | yes |
| L35 | 4 | 2 | L26 | | no |
| T37 | 9 | 2 | T28 | | no |
| F38 | 9 | 2 | F29 | | no |
| N41 | 9 | 1 | N32 | | no |
| F53 | 9 | 1 | V44 | | no |
| A57 | 5 | 2 | N48 | | no |
| F58 | 9 | 3 | V49 | | no |
| E60 | 8 | 4 | T52 | | no |
| F61 | 9 | 3 | L53 | | no |
| I62 | 8 | 1 | L54 | | no |
| G64 | 8 | 2 | S56 | | no |
| T65 | 8 | 3 | T57 | | no |
| L67 | 9 | 1 | L59 | | no |
| F68 | 9 | 2 | F60 | | no |
| F69 | 9 | 2 | L61 | | no |
| R71 | 9 | 2 | R63A | Glycoscore ~60% | no |
| D72 | 9 | 3 | E64 | | no |
| R75 | 9 | 3 | R67C | Glycoscore ~55% | yes |
| L124 | 7 | 3 | L92 | | no |
| L128 | 6 | 2 | F99 | | no |
| W131 | 4 | 2 | W102 | | no |
| R179 | 9 | 1 | K152E | Glycoscore ~60% | yes |
| E183 | 9 | 4 | E156 | | no |
| Y293 | 8 | 2 | F247 | | no |
| K299 | 9 | 3 | K253 | | no |
| L301 | 7 | 1 | I255 | | no |
| L302 | 9 | 1 | L256 | | no |
| T303 | 9 | 5 | T257 | | no |
| E304 | 9 | 3 | E258 | | no |
| D306 | 9 | 2 | E260 | Glycoscore ~60% | no |
| K307 | 9 | 5 | R261 | | no |
| L308 | 8 | 3 | Y262 | | no |
| N311 | 9 | 2 | 265 | | no |
| Y322 | 9 | 1 | Y278 | | no |
| A323 | 9 | 3 | D279 | | no |
| S326 | 8 | 5 | N282 | | no |
| N327 | 9 | 5 | N283 | | no |

Table S2

| | | | | | |
|---|---|---|---|---|---|
| Y328 | 9 | 1 | L284 | | no |
| S330 | 9 | 5 | S286 | | no |
| L331 | 8 | 3 | L287 | | no |
| L332 | 7 | 1 | V288 | | no |
| T333 | 9 | 5 | A289 | | no |
| R334 | 9 | 5 | R290A | Loss of function | no |
| F337 | 9 | 3 | F293 | | no |
| A338 | 7 | 3 | N294 | | no |
| E341 | 9 | 4 | E297 | | no |
| E342 | 9 | 2 | E298 | | yes |
| I381 | 9 | 1 | I343 | | no |
| G385 | 9 | 1 | G347 | | no |
| S389 | 8 | 1 | S351 | | no |
| L392 | 8 | 2 | A354 | | no |
| L396 | 6 | 2 | Y358 | | no |
| I410 | 8 | 1 | L372 | | no |
| Y413 | 9 | 1 | Y375 | | no |
| C414 | 7 | 2 | C376 | | no |
| Y416 | 9 | 2 | Y378 | | no |
| I417 | 7 | 1 | V379 | | no |
| L420 | 9 | 5 | L382 | | no |
| S421 | 8 | 1 | A383 | | no |
| G424 | 9 | 4 | G386 | | no |
| M479 | 9 | 3 | M439 | | no |
| R482 | 9 | 1 | R442A | Loss of function | yes |

* Number of times the residues are within 5Å of M5-DLO in the predicted models

^ For those residues that were mutated and tested, the mutation is indicated. Tests were done by using plasmid shuffling to introduce the variant of interest into RFT1-null yeast cells. Expression was driven by the strong GPD promoter. For phenotypes indicated as 'Loss of function' no viable cells were recovered after plasmid shuffling. For variants with the ability to support cell growth, CPY glycosylation was assessed. The CPY Glycoscore for cells expressing wild-type human Rft1 is ~70%. Data are taken from Hirata et al. (2024) J. Biol. Chem. 300: 107584 (PMID: 39025454).
